# Dynamics and Regulation of mRNA Cap Recognition by Human eIF4F

**DOI:** 10.1101/2025.06.26.660926

**Authors:** Hea Jin Hong, Matthew G. Guevara, Siyu Li, Yankang Liu, Alexandra N. Huang, Eric Lin, Arrmund Neal, Duo Xu, Rong Hai, Roya Zandi, Seán E. O’Leary

## Abstract

Efficient eukaryotic messenger RNA translation requires dynamic collaboration between the three subunits of initiation factor 4F (eIF4F, eIF4E•G•A), which recognises and activates mRNA at its 5ʹ cap structure for ribosome recruitment. Despite its high biological and pharmacological importance, the dynamics of full human eIF4F–mRNA engagement remain largely uncharacterised, hindering mechanistic understanding of translation initiation and its regulation. Here we observed human eIF4F activity with single-molecule fluorescence assays that directly visualise mRNA cap recognition by its eIF4E subunit. Unexpectedly, we find that inherently transient eIF4E–cap binding is repressed by full-length human eIF4G, predominantly through its C-terminus, representing an unanticipated role for eIF4G as a central rate-limiting factor in the eIF4F complex. This repression is relieved by nucleotide-bound eIF4A in the eIF4F heterotrimer, placing eIF4A as a crucial determinant of efficient cap recognition for translation. Molecular dynamics simulations reveal that electrostatic modulation of eIF4E–mRNA interaction allows eIF4G to control the cap-recognition frequency. Our findings also indicate that intrinsic eIF4F– mRNA dynamics are insufficient to support cap-tethered ribosomal scanning to locate translation start sites. They illuminate fundamental design-principle differences for the overall mechanism and division of labour among eIF4F subunits during mRNA recognition in humans and yeast.

## INTRODUCTION

mRNA translation is a central process in gene expression. For most eukaryotic mRNAs, efficient translation depends on recognition of their 5ʹ-terminal cap structure, m^7^G(5ʹ)ppp(5ʹ)N, where N is the +1 transcript nucleotide. The cap and its interacting proteins form a key cellular regulatory node,^1–6^ and enable efficient translational responses to extracellular stimuli.^7^ Dysregulated cap recognition is integral to cancer, viral infection, and developmental disorders,^8–10^ and is thus an attractive therapeutic target.^11–13^

The cap is recognised by eukaryotic translation initiation factor 4F (eIF4F), which is composed of eIF4E, eIF4G, and eIF4A protein subunits. Among these, eIF4E directly binds the cap structure,^14–17^ an interaction canonically regarded as the landmark initial recognition event preceding full eIF4F–mRNA engagement that facilitates ribosome recruitment.^18–20^ eIF4F–mRNA interaction is inherently dynamic, involving formation and reshaping of multiple protein–RNA contacts through eIF4G and eIF4A,^14,21–24^ and RNA structure remodeling by eIF4A helicase activity.^22–27^ Concerted, multivalent, and time-dependent eIF4F subunit activities during mRNA recognition make experiments to understand its molecular mechanism highly challenging, especially in the physiologically-critical context of full-length mRNA molecules.

Notably, no study has directly quantified how coordinated eIF4F subunit activities dictate mRNA binding and release dynamics for full human eIF4F on the translation initiation timescale. The current molecular-mechanistic model for human eIF4F function derives from studies of eIF4E or eIF4F subcomplexes containing eIF4G fragments, interacting with m^7^G(5ʹ)ppp(5ʹ) or short capped or uncapped olignucleotides.^28–30^ Critical questions thus remain unanswered about human eIF4F function and regulation.

A central challenge in understanding translation initiation is capturing molecular events on their native timescale, as these dynamics critically influence the efficiency and regulation of protein synthesis. Yet, even at this fundamental level, key questions remain unanswered: how often does human eIF4E locate mRNA cap structures, and how long does the interaction persist once formed? Answering these questions is essential for distinguishing between contrasting mechanistic models of initiation. For instance, whether the 5ʹ end of eIF4F-bound mRNA enters the ribosome *via* “threading” or “slotting” influences how 5ʹ-proximal start codons are selected.^31–33^ Similarly, whether eIF4F remains attached to the cap during ribosomal scanning to locate the translation start site, making the process “cap-tethered” or “cap-severed”, dictates how frequently initiation events can occur.^34–36^ More broadly, why different mRNAs vary in their sensitivity to eIF4F-mediated regulation and dysregulation^37,38^ remains incompletely understood. These insights have far-reaching implications for targeting translation initiation in disease intervention.

Recent single-molecule kinetic data for yeast eIF4F and past equilibrium-binding data for mammalian eIF4F challenge the eIF4E-centered mRNA recognition model,^39,40^ Instead, they place eIF4G– and eIF4A–mRNA interactions as preceding eIF4E–cap recognition, though the kinetic pathway remains uncharacterised for human eIF4F interactions with full-length mRNAs, and the relative importance of eIF4G and eIF4A in yeast and humans remains to be determined.^14^

Single-molecule approaches robustly parse component activities in concerted dynamic systems such as the eIF4F–mRNA complex. Building on our experimental approach for yeast eIF4F,^41^ we developed a fluorescently-labeled human eIF4E reagent to directly observe eIF4E and eIF4F real-time interactions with single human mRNAs. Our system includes full-length human eIF4G purified from human cells, which has not been utilized in past kinetic studies to our knowledge. With this system, we illuminate the kinetic basis of human eIF4F–mRNA recognition, including the contributions of eIF4G, eIF4A, and ATP to mRNA recognition.

Our study combined single-molecule analyses with sophisticated molecular dynamics simulations, enabling us to examine how mRNA structure and electrostatic forces guide eIF4F–mRNA recognition, which revealed a remarkably close alignment between our theoretical models and experimental findings. This agreement emphasizes the important role of electrostatic interactions in the system. More broadly, our findings underscore how functionally-conserved translation factors employ diverse molecular strategies to achieve efficient cap recognition in different organisms. By providing clear, interpretable insights into the kinetics of cap recognition, we propose a model of eIF4F–mRNA recognition that advances our understanding of translational control and the regulation of gene expression.

## RESULTS

To directly observe human eIF4E interaction with the mRNA cap in real time, we adapted our single-molecule fluorescence resonance energy transfer (smFRET) assay previously developed for yeast eIF4E.^41,42^ We fluorescently labeled human eIF4E at a genetically-encoded *p*-azidophenylalanine residue, by reaction with a Cy5 alkyne (Extended Data Figure 1a,b).^43,44^ Cy5-eIF4E includes the full native sequence extended by an N-terminal octapeptide, GHMA(*pAz*-Cy5)FMLE. Preservation of physiological eIF4E function even with N-terminal extension by polypeptides as large as the HaloTag (∼33 kDa) has been extensively validated *in vivo*.^45^

To quantify eIF4E–mRNA interaction frequencies and durations, RNAs were captured with biotin-5ʹ-(dT)_45_-3ʹ-Cy3 through their poly(A) tails, then immobilized on biotinylated zero-mode waveguide (ZMW) surfaces pre-treated with NeutrAvidin (Figure 1a).^41^ Cy5-eIF4E–mRNA interactions, reported by smFRET, were observed with data acquisition at 10 frames per second.^41,46^ During 532-nm Cy3-RNA excitation, eIF4E–cap binding brings the protein- and RNA-bound fluorophores into close proximity in the illumination volume, permitting smFRET from Cy3 to Cy5. eIF4E–mRNA binding at the cap therefore results in an instantaneous decrease in Cy3 fluorescence concomitant with the appearance of Cy5 fluorescence (Figure 1b), even though Cy5 is not directly excited. The end of each smFRET event marks eIF4E dissociation. smFRET events were essentially abolished by 1 mM m^7^GpppG dinucleotide, a competitive inhibitor of eIF4E–cap binding, confirming they specifically report on eIF4E binding to the mRNA cap (Extended Data Figure 1c).

**Figure 1.**
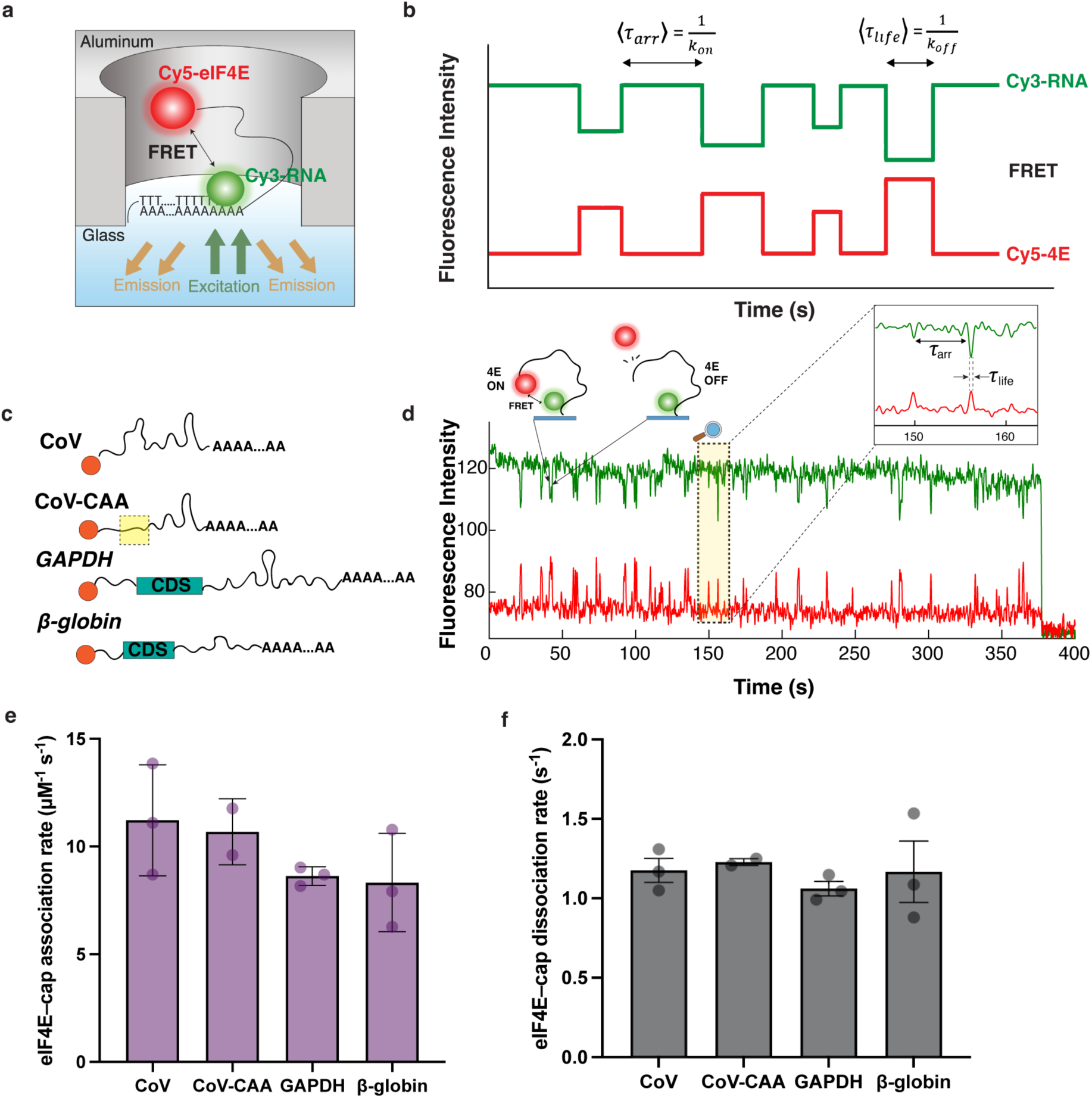
Single-molecule assay measuring eIF4E–cap interaction. **a.** Schematic of experimental design showing immobilization strategy and the basis of the smFRET signal for eIF4E–cap binding in a zero-mode waveguide. **b.** Schematic idealized single molecule fluorescence trace, showing the FRET signal obtained upon eIF4E–cap binding. The mean waiting time between FRET events provides the eIF4E association rate and mean duration of FRET events provides the dissociation rate. **c.** Schematic of the RNAs used in the single-molecule experiment. **d.** Representative single-molecule fluorescence trace for eIF4E interaction with the surface-immobilized SARS-CoV-2 5’ leader **e.** Second-order rate constants calculated for association of eIF4E with the viral leader (CoV), viral leader with CAA repeats in the 5′-proximal stem-loop (CoV-CAA), full-length human *GAPDH* and *β-*globin mRNAs. **f.** Rates of eIF4E–cap dissociation from the tested RNAs. Error bars in **e** and **f** and kinetic data throughout, reflect standard deviations from analysis of at least two independent experiments, each containing a minimum of 95 molecules.

This FRET-based assay reports eIF4E–cap engagement when mRNA ends are within ∼2 – 8 nm.^41^ Experimental and theoretical work indicate the ends of ∼400 – 2,000 nucleotide mRNAs lie within this distance,^47,48^ though the end-to-end distance distribution may be broad in thermally-accessible RNA conformational ensembles.^49^ To visualize eIF4E binding to all mRNA conformations, rather than only those permitting FRET, we implemented a second illumination scheme that directly excited both Cy3 and Cy5 during data acquisition. This revealed two dominant populations of mRNA molecules: those where the large majority of the eIF4E binding events resulted in FRET, and those where no events resulted in FRET (Extended Data Figure 1d,e). m^7^GpppG abolished the no-FRET binding events (Extended Data Figure 1f), confirming they also report on eIF4E–cap binding, but for mRNAs where the poly(A) tail is beyond ∼8 nm from cap-bound eIF4E. Notably, eIF4E interaction kinetics were indistinguishable between the sub-populations (Extended Data Figure 1g). Taken together, these assays enable robust quantitation of human eIF4E–mRNA interaction dynamics.

We compared eIF4E interaction dynamics between two full-length human mRNAs, β-globin and *GAPDH,* and the SARS-CoV-2 5ʹ untranslated region (“CoV”) (Figure 1c), a known and well defined sequence for initiating cap-dependent protein synthesis. Cap binding was invariably reversible and transient, broadly resembling yeast eIF4E (Figure 1d).^41,42,50^ Association rates (*k*_on_) were 8.6 ± 0.3 µM^−1^ s^−1^, 8.3 ± 1.3 µM^−1^ s^−1^, and 11.2 ± 1.2 µM^−1^ s^−1^ for *GAPDH*, β-globin and CoV, respectively (Figure 1e). They depended linearly on Cy5-eIF4E concentration up to ∼15 nM, confirming they report reliably on bimolecular eIF4E–mRNA binding in this range; we typically performed experiments at 7.5 nM (Extended Data Figure 2a). Binding was at least an order of magnitude slower than for short, capped RNA oligonucleotides,^51^ and slower than if driven by diffusional encounter.^52^ Slower-than-diffusion association rates are commonly found for RNA-protein interactions.^53,54^ Thus, the frequency with which human eIF4E locates and binds to the cap structure is significantly dictated by the mRNA “body”.

To probe effects of cap-proximal RNA structure, we leveraged the modular secondary structure of the SARS-CoV-2 UTR, where the cap-proximal stem-loop is structurally separable from the remainder of the RNA. We replaced the stem loop, which begins at the transcript +6 nucleotide, with an unstructured CAA-repeat sequence (“CoV-CAA”; Figure 1c); we validated *in silico* that the replacement represents a *bona fide* isolated structural change (Extended Data Figure 2b). eIF4E binding kinetics were unchanged within experimental error on CoV-CAA (*k*_on_ = 10.9 ± 0.9 µM^−1^ s^−1^; *k*_off_ = 1.2 ± 0.1 s^−1^) relative to native CoV RNA (Figure 1e). This contrasts with yeast eIF4E, which showed high sensitivity to cap-proximal structure in similar assays,^50^ but echoes a past report that inhibitory effects of cap-proximal secondary structures on translation diminish when the structures are positioned nine or more nucleotides from the cap. ^51^

We developed an analytical calculation to gain deeper insights into the principles underlying the differences between yeast and human eIF4E–mRNA association kinetics (Extended Data Figure 3a; Supplementary Note). The calculation is based on a diffusion-reaction model that takes account of (1) the diffusion of eIF4E particles outside the RNA’s radius of gyration, and (2) their diffusion within the RNA’s radius of gyration, where the potential of mean force becomes significant. This potential incorporates a combination of both specific and non-specific interactions between eIF4E and RNA, such as depletion and electrostatic forces, excluded volume effects, and other specific contacts. It reflects the overall strength of eIF4E–RNA interactions as eIF4E approaches the mRNA and surveys it to locate the cap structure. The calculation demonstrates that this potential between eIF4E and RNA dominates the length dependence of the association rate constant, *k*_on_. Stronger attractive eIF4E–RNA interaction results in a more pronounced dependence of *k*_on_ on RNA length (Extended Data Figure 3b). This framework provides a possible explanation for the nearly flat length dependence of *k*_on_ observed for human eIF4E (Figure 1e), in contrast to the stronger length dependence previously reported for the yeast factor.^41^

eIF4E dissociation rates were similar between mRNAs (*k*_off_ = 1.1 ± 0.1 s^−1^, *GAPDH* and CoV; 1.2 ± 0.1 s^−1^ CoV-CAA; 1.2 ± 0.2 *β*-globin) (Figure 1f), and up to three times faster than for yeast eIF4E, which exhibits a broader, mRNA-dependent distribution of dissociation rates.^41^ However, this eIF4E–mRNA binding duration was two orders of magnitude longer in our assay than reported for the dinucleotide cap structure or short capped oligoribonucleotides (∼10 ms)^30^; we did not observe highly transient binding even with data acquisition fast enough to unambiguously detect such events (i.e., with 12.5-milisecond movie frames) (Extended Data Figure 2c-e). Overall, our data demonstrate that while intrinsic eIF4E–mRNA interaction dynamics may allow eIF4E–cap binding to persist during mRNA loading into the ribosome, the intrinsic dynamics by themselves are insufficient to maintain eIF4F–cap contact throughout the seconds’ timescale of ribosomal scanning.^55–59^

In the prevailing mechanistic model, eIF4G promotes mRNA recognition for translation initiation by enhancing eIF4E–cap binding, largely through direct eIF4G–RNA binding.^42,60–65^ Kinetically, this enhancement could result from faster eIF4E•eIF4G–mRNA binding than for eIF4E alone, or from prolongation of the eIF4E•eIF4G•mRNA complex lifetime. Higher frequency or persistence of mRNA recognition increases the likelihood of subsequent ribosome recruitment. Indeed, yeast eIF4G enhances eIF4E–mRNA binding through both mechanisms.^39,41^ The eIF4E•eIF4G–mRNA binding duration is also a critical parameter in distinguishing between cap-tethered and cap-severed scanning mechanisms. However, the timing of human eIF4E•eIF4G engagement with full-length mRNAs remains unknown, in part because preparation of homogeneous, full-length human eIF4G is highly challenging.

We developed a method to purify homogeneous, full-length human eIF4G1 (residues 84-1600; Figure 2a) from human cells, based on transient overexpression followed by dual His_6_/FLAG affinity purification (Extended Data Figure 4a). Affinity tags at both protein termini allowed selective isolation of the full-length protein. Following past practice, we term eIF4G(84-1600) “full-length eIF4G”, as the N-terminal 83 residues are thought to be dispensable for function, and including them was reported to produce insoluble protein.^65,66^ We validated that purified full-length eIF4G stimulated eIF4A ATPase activity, using an NADH-coupled spectrophotometric assay (Extended Data Figure 4b). To investigate the role of eIF4G in eIF4E–cap binding, smFRET events for eIF4E–mRNA binding were observed in the presence of eIF4G(84-1600) at a concentration (25 nM) that saturates the eIF4E–eIF4G interaction (Figure 2b).^67^

**Figure 2.**
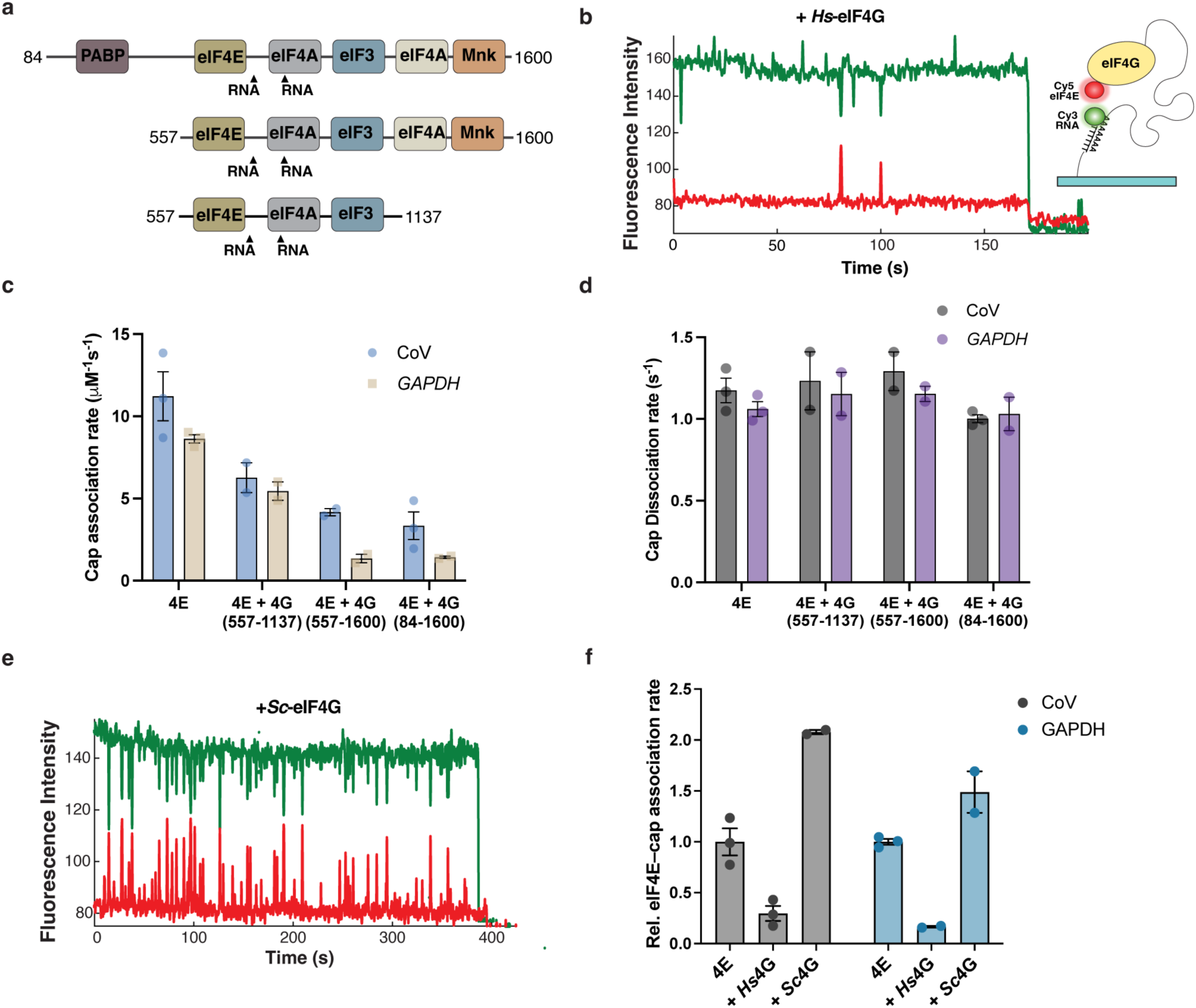
Effect of human eIF4G on eIF4E–cap interactions. **a.** Domain composition of human eIF4G proteins used for kinetic studies. **b.** Representative single-molecule fluorescence traces for eIF4E–cap interaction in the presence of full-length human eIF4G(84-1600) on CoV RNA and a schematic for the experimental setup **c.** eIF4E–cap association rates in the presence of eIF4G(557-1137, 557-1600, and 84-1600) on the CoV RNA (blue) and *GAPDH* (beige) mRNAs. **d.** eIF4E–cap dissociation rates in the presence of eIF4G(557-1137, 557-1600, and 84-1600) on the CoV RNA (grey) and *GAPDH* (purple) mRNA. **e**. Representative single-molecule fluorescence traces for eIF4E–cap interaction in the presence of yeast eIF4G on *GAPDH* mRNA. **f.** Relative eIF4E–cap association rate in the presence of full-length human or yeast eIF4G.

Unexpectedly, full-length eIF4G substantially decelerated eIF4E–mRNA association for CoV (*k*_on_ ∼ 3.4 ± 0.8 µM^−1^ s^−1^) and *GAPDH* mRNA (1.4 ± 0.1 µM^−1^ s^−1^) (Figure 2c), representing three-, four-, and six-fold decelerations. Moreover, in stark contrast to the yeast factor, human eIF4G did not prolong eIF4E–mRNA binding (Figure 2d). The aggregate effect was thus a three- to six-fold repression of eIF4E–cap recognition by eIF4G. Interestingly, deceleration was more pronounced with longer mRNAs, contrasting with the lack of a strong length dependence for eIF4E binding alone (Extended Data Figure 5a). Suppression of binding was not due to steric effects arising from the greater size of eIF4E•eIF4G relative to eIF4E, because full-length yeast eIF4G accelerated human eIF4E–cap binding to CoV (1.8-fold) and *GAPDH* mRNA (1.5-fold) (Figure 2e,f).

Taken together, the data indicate eIF4E attachment to the cap is intrinsically repressed with full-length human eIF4G1 in an mRNA length-dependent manner, which affects how frequently the cap is located by eIF4E•eIF4G, rather than how long the cap is engaged by eIF4E in the recognition complex. This suggests an unanticipated mechanism where eIF4G acts as a central rate-limiting factor for cap-recognition during initiation.

Repression of cap binding was largely a function of the eIF4G C-terminal segment. An eIF4G(557-1600) fragment (Figure 2a) behaved similarly to the full-length protein (*k*_on_: 4.2 ± 0.2 µM^−1^ s^−1^, CoV; 1.4 ± 0.3 µM^−1^ s^−1^, *GAPDH*). Conversely, an eIF4G(557-1137) fragment lacking the C-terminal segment (Figure 2a) decreased eIF4E–mRNA association rates by ∼40 % (*k*_on_ = 6.3 ± 0.9, CoV; 5.5 ± 0.6 µM^−1^ s^−1^, *GAPDH*) (Figure 2c). Again, dissociation rates remained unchanged for both fragments relative to full-length eIF4G, at ∼1 s^−1^ (Extended Data Figure 5b,c).

Recent smFRET kinetic data place yeast eIF4E•eIF4G-mRNA binding to the mRNA body through eIF4G as the most frequent initial point of eIF4E•eIF4G contact with mRNA.^39^ This echoes proposals, from a thermodynamic study of mammalian eIF4F, that eIF4G–mRNA interaction may generally precede eIF4E–cap interaction in the mRNA recognition mechanism.^40^ We tested this model for human eIF4E•eIF4G by again directly exciting both Cy5-eIF4E and Cy3-mRNA during data acquisition, to detect all interactions with mRNA, rather than only those that yield smFRET. As in the case of eIF4E alone, molecules with FRET and FRETless events were observed (Extended Data Figure 5d,e). However, the eIF4E–mRNA association rates determined by this approach were indistinguishable within experimental error from those measured by FRET alone (Extended Data Figure 5f). Moreover, under these direct Cy5 illumination conditions, movie acquisition at 80 fps did not reveal the presence of extra, highly-transient eIF4E•eIF4G–mRNA interactions that were initiated without FRET and then progressed to FRET due to repositioning of eIF4E•eIF4G from cap-distal to cap-proximal mRNA locations following a search process (Extended Data Figure 5g). Thus, if such a process occurs for human eIF4F, the time taken to complete it is extremely short (< 10 ms).

To probe mRNA-recognition dynamics of the full trimeric eIF4F complex, we supplemented smFRET reactions containing eIF4E (7.5 nM) and eIF4G (25 nM) with 1 µM eIF4A and 1 mM ATP·Mg (Figure 3a). These conditions were chosen to reflect the vast cellular excess of eIF4A over eIF4E•eIF4G.^68,69^ eIF4E–RNA smFRET was again observed in the presence of eIF4G and eIF4A (Figure 3b). Forming the eIF4F complex enhanced association rates across RNAs (*k*_on_ = 5.5 ± 1.0 µM^−1^ s^−1^, *GAPDH*; 5.3 ± 1.2 µM^−1^ s^−1^, CoV Figure 3c). Dissociation rates again remained unchanged relative to eIF4E•eIF4G, at ∼1 s^−1^ (Figure 3d). eIF4A in the eIF4F complex thus alleviates repression of eIF4E–cap binding by eIF4G solely by promoting the bimolecular association reaction.

**Figure 3.**
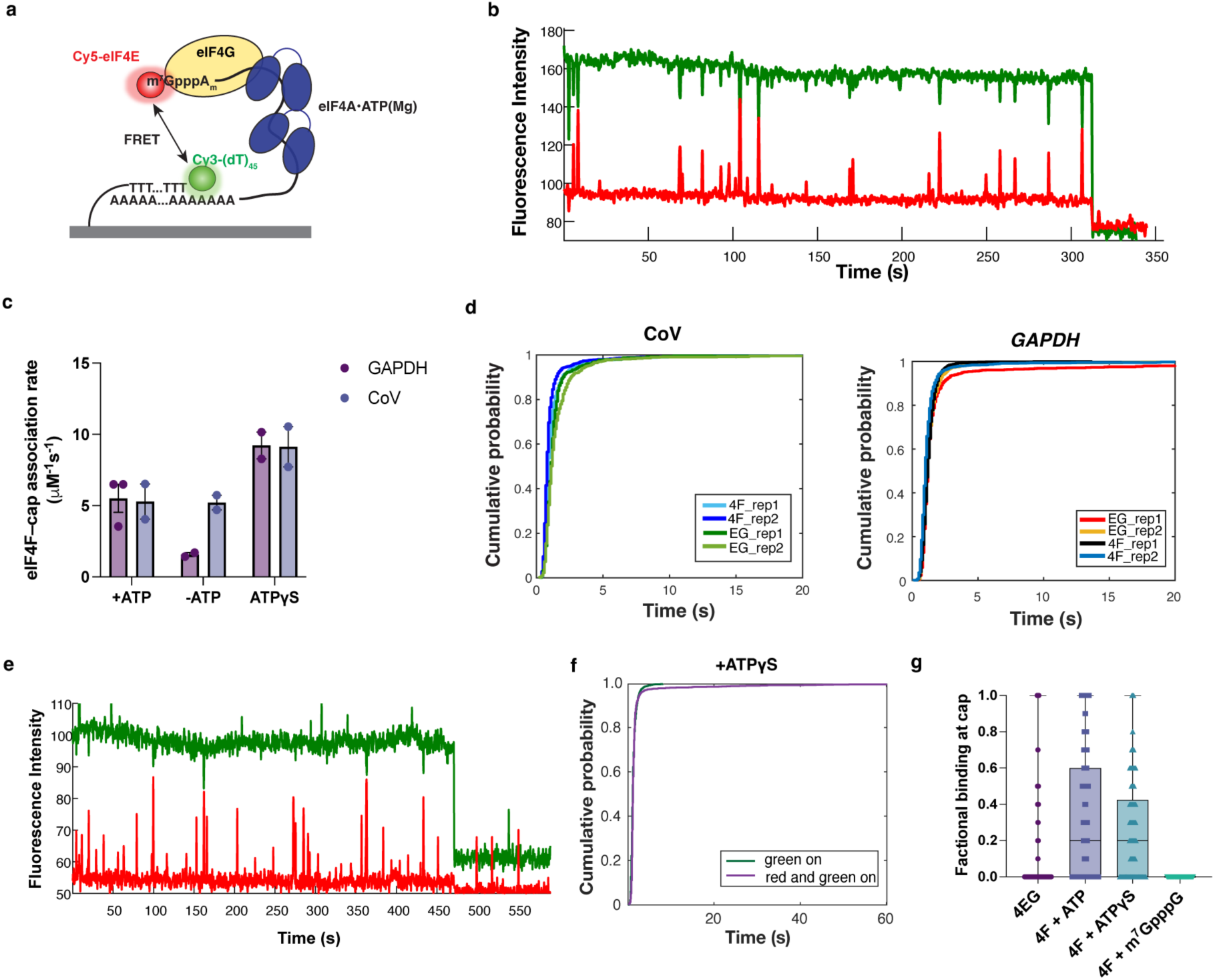
Effect of human eIF4A on eIF4EG–cap interactions. **a.** Schematic of experimental design showing eIF4E–cap binding within the full eIF4F complex. **b.** Representative single-molecule fluorescence trace for eIF4E–cap interaction in presence of human eIF4G and eIF4A. **c.** eIF4E–cap association rates in the presence of the full eIF4F complex, with ATP or ATP*γ*S, and without ATP, measured on the viral UTR or *GAPDH* mRNA. **d.** Cumulative distribution function of eIF4E–cap dwell times in the presence of eIF4G, or full eIF4F complex in two independent replicates. **e.** Representative single-molecule fluorescence trace for eIF4F–mRNA association on *GAPDH* mRNA in the presence of ATP*γ*S, with direct illumination of Cy5-eIF4E. **f.** cumulative distribution function of eIF4F dwell times on *GAPDH* mRNA **g.** Fraction of eIF4E–mRNA binding events occurring with FRET, representing cap binding, in the presence of ATP, ATP*γ*S, and m^7^GpppG.

Several biochemical activities could underpin this effect. Among the possibilities, eIF4A helicase or ATPase activity was not responsible. Instead, substituting ATP with the slowly-hydrolysable analogue, adenosine-5’-(γ-thio)triphosphate (ATPγS) almost doubled the eIF4F–mRNA binding rate relative to the ATP condition (*k*_on_,_ATPγS_ = 9.2 ± 0.9 µM^−1^ s^−1^, *GAPDH*; *k*_on_ = 9.1 ± 1.4 µM^−1^ s^−1^, CoV) (Figure 3c). This demonstrates that nucleotide binding was sufficient for eIF4A-enhanced cap recognition of *GAPDH* mRNA (*k*_on_,_–ATP_ = 1.6 ± 0.1 *μ*M^-1^ s^-1^; Figure 3c). Interestingly, though, the CoV UTR was able to efficiently bind eIF4F even in the absence of any nucleotide (*k*_on,–ATP_ = 5.2 ± 0.5 *μ*M^-1^ s^-1^; Figure 3c), possibly reflecting a viral strategy for efficient eIF4F utilisation.

A potential “tethering” mechanism for eIF4A promotion of eIF4F–mRNA recruitment relies on coupling between eIF4A-RNA and eIF4A-ATP binding.^70^ Here, eIF4A is the primary eIF4F contact point with mRNA. For such a mechanism, imaging eIF4F mRNA-recognition dynamics with direct Cy5-eIF4E excitation would yield a diagnostic signal: appearance of Cy5 fluorescence upon eIF4F–mRNA binding through eIF4A would be followed by transient excursions to FRET as eIF4E bound and released the cap. Since ATP hydrolysis is required for eIF4A release from RNA, its timescale (*E*_0_/*V* ∼100 s for the CoV RNA, similar to past reports^71^; Extended Data Figure 4) places a lower limit on the overall Cy5 event duration in this model, and ATPγS is expected to prolong that duration. However, all eIF4F–RNA binding events that we detected directly through Cy5-eIF4E remained transient, and order-of-magnitude shorter than eIF4A ATP hydrolysis, regardless of the presence of ATP or ATPγS (Figure 3e,f). Thus, the data do not support the tethering model.

The remaining possibility is that incorporation of eIF4A in eIF4F allosterically enhances eIF4E– cap binding. Indeed, allostery is integral to eIF4F function: eIF4E–eIF4G interaction activates the helicase activity of eIF4G-bound eIF4A,^72^ and eIF4E and ATP drastically enhance the affinity of eIF4A•eIF4G complexes for oligoribonucleotides.^21^ In the allosteric mechanism, nucleotide-bound eIF4A stabilises an “open” eIF4F conformation that is permissive for eIF4E–cap binding, reciprocating allosteric activation of eIF4A helicase activity by eIF4E. The cap-binding kinetics of eIF4F(ATP) across the mRNAs tested most closely resembled those for eIF4E•eIF4G(557– 1137), with common association rates of ∼5 – 6 µM^−1^ s^−1^, in contrast to the situation with eIF4E•eIF4G where the rates were lower and mRNA-specific.

The simplest explanation for these data is that inclusion of eIF4A(ATP·Mg) in the eIF4F complex relieved eIF4G C-terminal inhibition of eIF4E–mRNA recognition. Accordingly, the proportion of eIF4F–mRNA encounters that resulted in eIF4E–mRNA FRET increased from close to zero for the eIF4E•eIF4G complex to as much as ∼40-60% when ATP or ATPγS was present and returned to essentially zero when eIF4F(ATP·Mg) bound RNA in the presence of the cap-structure analogue m^7^GpppG (Figure 3g). Taken together, our data are consistent with a model where direct human eIF4E–cap binding becomes the fastest of the possible processes for mRNA encounter in the eIF4F complex, and thus kinetically controls mRNA recognition.

To deepen our understanding of the interplay between RNA properties and eIF4F subunits in cap recognition, we created a quantitative *in-silico* molecular dynamics simulation model for eIF4E– and eIF4E•eIF4G–mRNA interaction with full-length mRNAs (Figure 4a, 4b; Extended Data Figure 6). We designed a coarse-grained eIF4E protein model based on the three-dimensional structure of murine eIF4E (98% sequence identity to human eIF4E).^73^ In this model, a region of eIF4E (purple; Figure 4a) interacts attractively with the cap particle (green), while a positively charged site at the center of the protein (Extended Data Figure 6) interacts electrostatically with the negatively charged mRNA and cap.

**Figure 4.**
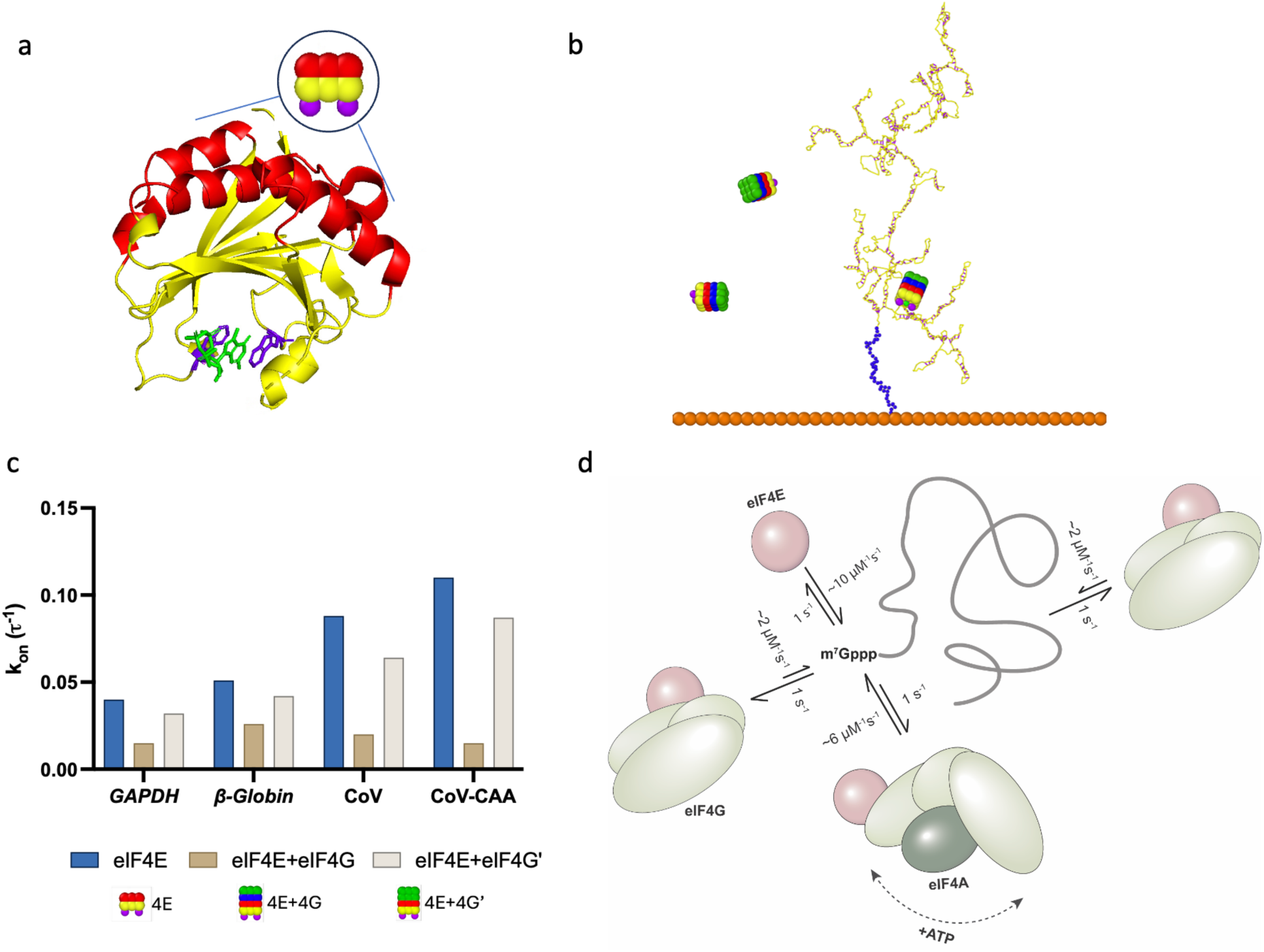
Molecular dynamics simulations reveal that eIF4G represses eIF4E-cap binding. **a**. Illustration of the eIF4E protein structure and coarse-grained design. The two tryptophan residues are coloured in purple and the mRNA cap structure is coloured in green. **b.** Molecular dynamics snapshot illustrating one surface-tethered *GAPDH* RNA surrounded by eIF4E and eIF4G proteins. **c.** Simulated association rates of eIF4E, eIF4E+eIF4G, and eIF4E+eIF4G’ complexes binding to mRNAs with varying lengths and secondary structures. **d.** Allosteric model of eIF4F cap–binding.

Four different mRNAs, corresponding to those studied in the single-molecule assays, were examined with these simulations: *β*-globin, *GAPDH*, the SARS-CoV-2 viral leader sequence, and its variant where stem-loop 1 is replaced by CAA repeats. In the simulations, we assumed that the mRNA molecules adopted their minimum-energy secondary structures as predicted by the ViennaRNA package (Extended Data Figure 2b).^74^ For each simulation, the cap particle is attached to the mRNA 5ʹ end and a linear poly(A) chain to the mRNA 3ʹ end, which in turn is anchored to a surface (orange) to recapitulate mRNA immobilization in the single-molecule fluorescence experiments. An example of the simulation is shown in Figure 4b, where three proteins diffuse around the RNA. Of these, one binds to the cap. The binding event is registered when the centre of the protein comes within 5 nm of the mRNA 3ʹ end, echoing the experimental detection of smFRET.

The simulation was conducted in a cubic box of dimensions 50 nm × 50 nm × 50 nm containing five proteins, corresponding to an effective concentration of approximately 66 µM. Although this is higher than experimental conditions, it preserves the relative ratios of molecular species to reflect the microenvironment within fluorescence-positive wells. These wells, which contain one or more captured molecules, exhibit higher local concentrations due to the stochastic nature of molecular loading across the full well array.

The simulated eIF4E–mRNA association rates (*k*_on_) were 0.040 τ^−1^, 0.051 τ^−1^, 0.088 τ^−1^, and 0.11 τ^−1^ for *GAPDH*, *β*-globin, CoV, and CoV-CAA, respectively (Figure 4c), where τ is the unit time of the simulations. Dissociation rates were similar across mRNAs, with *k*_off_ = 0.44 τ^−1^for GAPDH, 0.41 τ^−1^for *β*-globin; 0.37 τ^−1^ for CoV, and 0.37 τ^−1^ for CAA–CoV-2. While the association rates vary across the different RNA targets, the dissociation rates are all on the same order of magnitude. Thus, the simulated binding dynamics are in good agreement with the experimental behaviour of eIF4E.

We applied this system to understand the properties of eIF4G that slow eIF4E–cap recognition in the eIF4E•eIF4G complex, by adding to the simulation an eIF4G particle irreversibly bound to eIF4E, to represent the eIF4E•eIF4G complex. We performed simulations that included two versions of eIF4G. In the first, the eIF4G particle was negatively charged in the layer connected to eIF4E, reflecting the expected physiological state of eIF4G, which has an overall isoelectric point (pI) of ∼5.2. The second version was neutral at this position, mimicking the eIF4G•eIF4A complex (eIF4G*ʹ*; Figure 4c), in which a negatively charged surface of eIF4G is occluded by eIF4A binding.

When negatively charged, eIF4G substantially reduced the eIF4E–mRNA association rate for all mRNAs, with *GAPDH* (*k*_on_ = 0.015 τ^−1^), β-globin (0.026 τ^−1^), CoV (0.020 τ^−1^) and CAA-CoV (0.015 τ^−1^). However, when the modified form of eIF4G with a neutral interaction layer was used, the association rates were partially restored, with *GAPDH* (0.032 τ^−1^), β-globin (0.042 τ^−1^), CoV (0.064 τ^−1^) and CAA-CoV (0.087 τ^−1^). These data strongly support a role for the electrostatic properties of eIF4G, modulated by eIF4A, in controlling eIF4E–cap recognition beyond any specific RNA-binding domain.

## DISCUSSION

Collectively, our data provide new insights into the dynamics underpinning mRNA recognition for human translation, its preparation for ribosomal pre-initiation complex (PIC) recruitment, and the fundamental dynamics of eIF4F that occur on the scanning timescale. The transient nature of intrinsic eIF4E–cap engagement, which is invariant with mRNA identity, places the eIF4E–cap bimolecular association rate as a crucial determinant of mRNA recognition, consistent with kinetic control of the initiation process. Our simulations and theoretical calculations underscore the importance of electrostatics in eIF4F–mRNA recognition, highlighting their importance in controlling a protein’s search for a small binding site on a large mRNA molecule.

The transient eIF4E–cap binding we observe, even for the full eIF4F•mRNA complex, places important constraints on the mechanistic requirements for PIC recruitment and scanning. Since we find that eIF4G or eIF4F do not prolong eIF4E–cap engagement, if scanning is cap-tethered in human cells, the tethering may require additional mRNA contacts beyond the eIF4F core. Potentially stabilising interactions arising from eIF4G–mRNA anchoring by the poly(A)-binding protein might represent one such avenue for tethering, consistent with the “closed loop” model for translation initiation.^4^ Alternatively, mRNA binding by other PIC-associated factors may be involved, such as eIF3.^33^

Our study highlights how the dynamic properties of human eIF4F differ significantly from its yeast counterpart in several important aspects. In both systems, eIF4G clearly plays a defining role in driving cap recognition. In yeast, this is through lengthening eIF4E•eIF4G–RNA encounters to promote cap recognition in the correct context.^39,41^ In the human system, eIF4G evidently acts instead as a “brake” on eIF4E–cap engagement outside the full eIF4F complex. Our findings are also consistent with and extend the “autoinhibition” model proposed for eIF4G function, where eIF4A activity in the eIF4G•eIF4A complex is inherently suppressed, but this suppression is relieved by eIF4E binding.^72^ Here we show that this autoregulation is reciprocal – eIF4A–eIF4G binding also activates eIF4E (Figure 4d). Ultimately, though, in both systems it is nucleotide-bound eIF4A that sets the eventual efficiency of forming a cap-bound eIF4F•mRNA complex (Figure 4d).

Overall, our findings for human eIF4F align with a more protein-centric regulatory model for the human system than is found for yeast. Rapid recycling of eIF4F–mRNA interactions on the initiation timescale, and the associated importance of kinetic control of cap recognition, magnifies the impact of post-translational modifications or other regulatory mechanisms that impact the ability of eIF4F to find mRNA cap structures. This is consistent with the additional layers of levels of extrinsic, protein-directed regulation found for human eIF4F relative to the yeast factor, such as eIF4E serine-209 phosphorylation, which plays an important role in translation regulation. Future studies to characterise the effects of regulatory processes on real-time eIF4F–mRNA dynamics will deepen our understanding of this central process in gene expression.

## MATERIALS AND METHODS

### Cloning and synthesis of RNA constructs

DNA templates for *in vitro* RNA transcription of SARS-CoV-2 RNAs were obtained commercially (Azenta, Inc.). For preparation of the viral UTR construct, DNA fragments containing the class II T7 promoter Φ2.5, followed by the SARS-CoV-2 5’ untranslated region sequence were assembled in a pUC57 vector. An EcoRI restriction site was placed at the 3’ end of the insert, to allow template linearization prior to transcription. For CoV-CAA, the first 5’ proximal stemloop of SARS-CoV-2 UTR construct was replaced with 6 CAA repeats. The full-length human *GAPDH* transcript sequence, obtained from UCSC Genome Browser, was purchased as a gBlock from IDT. The sequence was inserted into pUC119 *via* SalI and EcoRI restriction sites incorportated during gBlock synthesis. A T7 promoter sequence was also included at the 5’ end, along with an extra 5’ dG to facilitate efficient *in vitro* transcription. The EcoRI site placed at the 3’ end allowed linearization of the template for the subsequent transcription reaction. The full-length human *β*-globin transcript sequence was purchased as a gBlock from IDT. The sequence was inserted into pUC119 *via* HindIII and SalI restriction sites incorportated during gBlock synthesis. A T7 promoter sequence was also included at the 5’ end, along with an extra 5’ dG to facilitate efficient *in vitro* transcription. The SalI site placed at the 3’ end allowed linearization of the template for the subsequent transcription reaction. Plasmids were isolated at preparative scale from *E*. coli DH5α transformants selected with ampicillin (100 µg/mL), using a maxi-prep kit (Macherey Nagel). The insert DNA sequences for each construct are given in Supplementary Information. The sequences of all plasmid inserts used in this study were confirmed by Sanger sequencing (Azenta, Inc.).

### RNA transcription

Purified plasmids were digested at the 3’ end of the insert sequence with EcoRI-HF (NEB) to linearize the template with a 5’ overhang for *in vitro* transcription. For 60 µL transcription reactions, about 11 µg of linearized DNA template was incubated for 4 h at 37 °C with 12.5 mM of each NTP, 3% (v/v) DMSO, 25 mM MgCl_2_, 17 mM DTT, and T7 RNA polymerase (12,500 units/mL) in transcription buffer composed of 0.4 M Tris-HCl (pH 8.1), 10 mM spermidine, and 0.01% (v/v) Triton X-100. The transcription product was extracted using acidic phenol chloroform (pH 4.5) and precipitated overnight in ethanol at –20 °C. RNA was redissolved in water. RNAs were capped using the *Vaccinia* capping system (NEB) and poly(A)-tailed at their 3’ ends with poly(A) polymerase (NEB), following the manufacturer’s protocol. RNA concentration was quantified by UV absorbance spectrophotometry using a NanoDrop instrument.

### Preparation of fluorescently-labeled human eIF4E

A pET-28a(+) vector encoding the human eIF4E sequence fused with an N-terminal Protein G tag, were designed similarly to Feokistova *et. al*. (2013).^72^ The construct contained a sequence encoding the peptide MA(*p*AzF) between the Protein G tag and the N-terminus of eIF4E, where *p*AzF is *p*-azidophenylalanine (pAzF). *p*AzF was encoded by the amber stop codon (TAG), for decoding by *p*AzF-tRNA^CUA^. The insert sequences are given in the Supplementary Information.

The pULTRA expression vector, containing inserts encoding tRNA^CUA^ and pAzF-tRNA^CUA^-synthetase – a generous gift from Abhishek Chatterjee, Boston College – was co-transformed into an *E. coli* BL95ΔAΔ*fabR* strain with the pET28a(+)-eIF4E plasmid, and transformants were selected on LB agar containing Kanamycin (50 µg/mL) and Spectinomycin (100 µg/mL). A single colony from this selection was used to inoculate a 10 mL 2×YT broth starter culture containing both antibiotics, which was grown overnight at 37 °C. The following day, the starter culture was used to inoculate 1 L of 2xYT broth containing the antibiotics. The culture was grown at 37 °C to OD_600_ ∼0.6, and then 1 mM pAzF (BAChem, F3075) was added along with 1 mM isopropyl-β-D-thiogalactopyranoside (IPTG) to induce recombinant protein overexpression. Induction was carried out for 5 h at 30 °C in darkness. The cells were spun down and suspended in lysis buffer (20 mM HEPES-KOH pH 7.5, 400 mM KCl, 5 mM imidazole, 1×EDTA-free protease inhibitor cocktail (0.24 mg/mL benzamidine hydrochloride hydrate, 2 µM pepstatin, and 0.6 µM leupeptin hemisulfate, 1× phenylmethylsulfonyl fluoride (PMSF)). Proteins were purified immediately after induction, without freezing of the cell pellet. Further purification was carried out on ice with ice-chilled buffers. The cells were lysed using a sonicator equipped with a microtip (Branson Sonifier 450; 80 % amplitude, setting 3, for 45 seconds at 5 min intervals) and the lysate was immediately clarified by centrifugation (22,000 rpm for 30 min). The filtered lysate was passed through 5 mL Ni-NTA agarose (Thermo Fisher), followed by typically 3 washes with 5 column volumes of wash buffer (20 mM HEPES-KOH pH 7.5, 400 mM KCl, 10 mM imidazole) until no further detectable protein eluted from the column, as assessed by Bradford reagent (Bio-Rad). Ni-NTA-bound protein was then eluted with elution buffer (20 mM HEPES-KOH pH 7.5, 400 mM KCl, 200 mM imidazole), and eluate fractions were analyzed by 10 % SDS-PAGE. Fractions that contained eIF4E were combined, and labeled overnight at 4 °C with 50 µM Cy5-DBCO (Sigma). The fluorophore was dissolved in 100% DMSO for addition to the labelling reaction; the final DMSO concentration in the labelling reaction was 1% (v/v). The labeled protein was desalted to remove unreacted dye and the sample was buffer exchanged into TEV cleavage buffer (20 mM HEPES-KOH, 50 mM KCl, 10 % (v/v) glycerol). The desalted sample was then incubated with TEV protease (NEB) (50 units) overnight at 4 °C for complete cleavage of protein G from the 4E. Once the cleavage was complete, the sample was loaded onto an SP-column (GE Healthcare Life Sciences) maintained at ∼4 °C. The column was washed with 20 column volumes of low-salt buffer (20 mM HEPES-KOH, 50 mM KCl, 10 % Glycerol), then eluted with a linear gradient from 0% to 100% high-salt buffer (20 mM HEPES-KOH, 500 mM KCl, 10 % Glycerol). Fractions containing Cy5-eIF4E typically eluted at ∼250 mM KCl. Based on inspection of an SDS-PAGE gel of the eluate fractions, imaged for Cy5 fluorescence by a Typhoon imager, Fractions containing labeled protein were combined, concentrated to < 1 mL by centrifugal ultrafiltration (10,000 or 5,000 MWCO ultrafilter,Cytiva) and loaded onto a Superdex 75 Increase (10/300 GL) column (GE Healthcare Life Sciences) equilibrated in storage buffer (20 mM HEPES-KOH pH 7.5, 200 mM KCl, 10 % Glycerol, 1 mM TCEP) for further purification. Fractions were assessed for purity by SDS-PAGE, and for labelling efficiency by measuring the ratio of 640 nm to 280 nm absorbance. The 1,2,3-triazole formed in the azide-alkyne cycloaddition reaction that conjugates the fluorophore to the protein is expected to absorb at 280 nm. Considering the contribution of this additional absorbance, the typical labelling efficiency was ∼50%; this assessment was supported by SDS-PAGE analysis of the labelled protein, which contained two bands, only one of which was conjugated to Cy5. Purified Cy5-eIF4E were stored at 4 °C in darkness and used for a maximum of two weeks after each purification.

### Preparation of human eIF4A

pHis APEX2-eIF4A1 was a gift from Nicholas Ingolia (Addgene plasmid #12964; Padrón et. al., 2019). The plasmid was transformed into BL21(DE3) CodonPlus cells and transformants were selected on LB–agar plates containing ampicillin (100 µg/mL) and chloramphenicol (25 µg/mL). A single colony from this selection was used to grow a 10 mL overnight culture at 37 °C. This starter culture was then used to inoculate 1 L LB containing the selective antibiotics. The cells were grown to OD_600_ of ∼0.6, then protein overexpression was induced with 1 mM IPTG for 3 h at 37 °C. After induction, the cells were harvested by centrifugation at 4,000 rpm for 15 min and were frozen and stored at -80 °C until purification. The frozen cell pellet was resuspended in lysis buffer (20 mM HEPES-KOH, pH 7.5, 500 mM NaCl, 10 mM imidazole, 10 mM β-mercaptoethanol, 0.5% (v/v) NP-40). The cells were lysed by sonication with a microtip attachment (80 % amplitude, setting 3, for 45 seconds at 5 min intervals) and the lysate was immediately clarified by centrifugation (22,000 rpm for 30 min). During centrifugation, a gravity-flow column containing 1 mL Ni-NTA agarose (Thermo Scientific) was washed and pre-equilibrated with the lysis buffer. The cell-free supernatant was diluted twofold with this lysis buffer and filtered through a 0.22 µm syringe filter (Corning). The filtrate was then applied to the equilibrated Ni-NTA column, which was washed with a further ∼20 mL lysis buffer. The column was further washed with 40 mL high-salt buffer (20 mM HEPES-KOH, pH 7.5, 1 M NaCl, 20 mM imidazole, 10 mM β-mercaptoethanol), followed by low-salt buffer (20 mM HEPES-KOH, pH 7.5, 500 mM NaCl, 20 mM imidazole, 10 mM β-mercaptoethanol). The protein was then eluted with elution buffer (50 mM Na-phosphate buffer, pH 7.5, 500 mM NaCl, 100 mM Na_2_SO_4_, 250 mM imidazole, 2 mM DTT). After elution, the eluate was buffer-exchanged into low-salt buffer (20 mM Tris HCl pH 7.5, 100 mM KCl, 5% glycerol, 2 mM DTT, 0.1 mM EDTA) using a 10 DG column (Bio-Rad), and was TEV-protease (50 units; NEB) cleaved overnight at 4°C to remove the APEX tag. The TEV-protease cleaved sample was loaded onto a 5 mL Q-Sepharose HP column (GE Healthcare Life Sciences), equilibrated in low-salt buffer and maintained at ∼4 °C. The column was washed with 20 column volumes of low-salt buffer, then eluted with a linear gradient from 0% to 100% high-salt buffer (20 mM Tris HCl pH 7.5, 500 mM KCl, 5% glycerol, 2 mM DTT, 0.1 mM EDTA) over 50 column volumes. eIF4A typically eluted at ∼265 mM KCl. Eluate fractions were analyzed by SDS-PAGE, and fractions containing pure eIF4A were then buffer exchanged into a storage buffer (20 mM Tris HCl, pH 7.5, 2 mM DTT, 0.1 mM EDTA, 10% glycerol, 100 mM KCl) using Superdex-75, flash frozen under liquid nitrogen and was stored at -80 °C.

### Preparation of His_6_-tagged human eIF4G(557–1137)

A pET-28a(+) vector was constructed with an insert encoding a hexahistidine tag followed by residues 557 to 1137 of human eIF4G1, codon-optimized for *E. coli*. The plasmid was transformed into *BL21 (DE3) CodonPlus* cells. Transformants were selected on LB-agar plates containing kanamycin (50 μg/mL) and chloramphenicol (25 μg/mL). A single colony was used to inoculate a 10 mL LB starter culture, which was grown overnight at 37 °C. The following day, the starter culture was used to inoculate nine 1 L LB cultures containing the selective antibiotics. Cells were grown at 37 °C to an OD₆₀₀ of ∼0.6, at which point protein overexpression was induced by the addition of 1 mM IPTG. Cultures were incubated overnight at 16 °C. Cells were harvested by centrifugation at 4,000 rpm for 15 minutes and stored at –80 °C until purification. Frozen cell pellets were resuspended in ∼45 mL of lysis buffer (20 mM HEPES-KOH pH 7.5, 400 mM KCl, 5 mM imidazole, 10% glycerol, 10 mM β-mercaptoethanol, 1× EDTA-free protease inhibitor [Roche], 1× PMSF) and lysed using a microtip sonicator (80% amplitude, setting 3, for 30 seconds at 2-minute intervals). Approximately 4 mL of Ni-NTA agarose (Thermo Scientific) was washed and pre-equilibrated with lysis buffer while the lysate was clarified by centrifugation at 20,000 rpm for 30 minutes. The clarified lysate was filtered first through a 5.0 μm syringe filter (Corning), then through a 0.80 μm filter. The filtered lysate was applied to the equilibrated Ni-NTA column and washed with ∼15 mL of lysis buffer, followed by 50 mL of wash buffer (20 mM HEPES-KOH pH 7.5, 400 mM KCl, 10 mM imidazole, 10% glycerol, 10 mM β-mercaptoethanol). Ni-NTA– bound proteins were eluted in 1.0 mL aliquots using elution buffer (20 mM HEPES-KOH pH 7.5, 200 mM KCl, 200 mM imidazole, 10% glycerol, 10 mM β-mercaptoethanol). Protein-containing fractions, as assessed by SDS-PAGE, were pooled and buffer exchanged to reduce the salt concentration to ∼66 mM using a 10-DG desalting column (Bio-Rad), pre-equilibrated in elution buffer lacking imidazole and DTT. The buffer-exchanged protein was loaded onto four pre-equilibrated 1 mL heparin columns. The protein was eluted using a linear 0–100% gradient from low-salt buffer (20 mM HEPES-KOH, pH 7.5, 100 mM KCl, 10% glycerol, 2 mM DTT) to high-salt buffer (20 mM HEPES-KOH pH 7.5, 1 M KCl, 10% glycerol, 2 mM DTT). Protein-containing fractions (∼450 mM KCl) were diluted with no-salt buffer and loaded onto four 1 mL Q columns. The protein was then eluted using the same buffer system as used for the heparin column. Eluted fractions were pooled and dialyzed (Thermo Scientific; Product No. 69552) into storage buffer (20 mM HEPES-KOH, pH 7.5, 200 mM KCl, 10% glycerol, 1 mM DTT), flash-frozen in liquid nitrogen, and stored at –80 °C until use.

### Preparation of human eIF4G(84-1600 and 557-1600)

The expression plasmid for N-terminal His_6_-tagged and C-terminal FLAG-tagged eIF4G (encoding amino acids 84-1600) was generated by cloning of the corresponding DNA fragment into the pCAGGS vector, a mammalian expression vector driven by the chicken beta actin promoter (a gift from the laboratory of Dr. Peter Palese Laboratory). The plasmid was transfected into HEK293T cells. Transfected cells were pelleted for 15 min at 4,000 rpm and stored at -80 °C until purification. Codon-optimized N-terminal-His_6_ tagged, and C-terminal FLAG tagged eIF4G (557-1600) sequence was inserted in the pET28a(+) plasmid and was transformed in BL21-codon plus cells (Agilent). Frozen cells were resuspended in lysis buffer (20 mM HEPES pH 7.5, 400 mM KCl, 5 mM Imidazole, 10 % Glycerol, 10 mM β-mercaptoethanol, 1 x protease inhibitor, 1 x PMSF, 0.1 % NP-40) and was lysed using a microtip sonicator as described above (80 % amplitude, setting 3, for 30 seconds in 2-minute intervals). A 5 mL Ni-NTA Chelating column (Cytiva) was charged with 0.1 M NiSO_4_ (∼5 mL), washed with water (∼30 mL) and pre-equilibrated with lysis buffer (∼30 mL) while the lysate was clarified by centrifugation (18,000 rpm for 30 min). The clarified lysate was filtered first through 0.45 µm syringe filter (Corning) then by a 0.22 µm filter. The filtered lysate was added to the equilibrated column and washed with ∼20 mL lysis buffer. The column was washed with 50 mL of wash buffer (20 mM HEPES-KOH pH 7.5, 400 mM KCl, 10 mM Imidazole, 10 % glycerol, 10 mM β-mercaptoethanol, 1 x protease inhibitor, 1 x PMSF, 0.1 % NP-40). The proteins were eluted in three 1 mL fractions using 100 mM imidazole elution buffer (20 mM HEPES-KOH pH 7.5, 400 mM KCl, 100 mM Imidazole, 10 % glycerol, 10 mM *β*-mercaptoethanol, 0.1 % NP-40), followed by seven 1 mL fractions using 500 mM imidazole elution buffer (20 mM HEPES-KOH pH 7.5, 400 mM KCl, 500 mM Imidazole, 10 % glycerol, 10 mM *β*-mercaptoethanol, 0.1 % NP-40). The eluted samples were analyzed using nanodrop and SDS-PAGE.

The 500 *μ*L elution fractions containing eIF4G were loaded to 40 *μ*L of magnetic FLAG resin (Thermo Fisher) pre-equilibrated with elution buffer without imidazole (20 mM HEPES-KOH pH 7.5, 400 mM KCl, 10 % glycerol, 10 mM *β*-mercaptoethanol, 0.1 % NP-40). The protein was incubated overnight at 4 °C with constant rocking. The following day, the protein–bound FLAG resin was washed three times with equilibration buffer. The protein was eluted with 75 *μ*L of FLAG elution buffer (20 mM HEPES pH 7.5, 400 mM KCl, 150 ug/mL FLAG peptide, 10 % glycerol, 10 mM *β*-mercaptoethanol, 0.1 % NP-40) by incubating for 30 min at room temperature with constant rocking and pipetting out the eluate. Another 75 *μ*L of FLAG elution buffer was added and incubated for 30 min to elute the rest of the bound proteins. The eluted protein was analyzed using SDS-PAGE and was buffer exchanged into storage buffer (20 mM HEPES-KOH pH 7.5, 200 mM KCl, 10 % glycerol using a 10 kDa mini-dialyzer (Thermo fisher). The protein sample was flash frozen with liquid nitrogen and stored at -80 °C. The full-length eIF4G is prone for degradation, so it was purified fresh every 3 months. The protein was analyzed on an SDS-PAGE to check for degradation. eIF4G (557-1600) was purified the same way as the full-length.

### eIF4A ATPase assay

An NADH-linked enzyme coupled assay^75^ was performed to measure the ATPase activity of eIF4A (Bradley et al., 2012). All reagents were freshly prepared, including the stocks of 15 mM NADH, 100 mM Mg•ATP (pH 7.0), Lactate dehydrogenase (LDH; 4000 U/mL; Sigma 427217.), 100 mM phosphoenolpyruvate (PEP; pH 7.0) and 1M DTT. A 5 x coupling assay cocktail was prepared in 1 x KMg75 buffer (20 mM HEPES-KOH pH 7.5, 75 mM KCl, 5 mM MgCl_2_ and 1 mM DTT), with the following components in final concentrations: 1 mM NADH, 100 U/mL LDH, 500 U/mL pyruvate kinase, 2.5 mM PEP. A separate tube containing 2 x coupling assay cocktail with or without proteins (eIF4A or eIF4G or both) in 1 x KMg75 buffer and another tube containing 2 mM ATP and 2 µM CoV RNA in 1 x KMg75 buffer were prepared. The solutions were mixed in a 100 µL quartz cuvette with 1 cm pathlength and immediately monitored using a UV spectrophotometer (Shimadzu) at 340 nm for 15 min, at 0.7-s time intervals. The time course of absorbance change was then fit by linear regression to obtain the slope. This and the background NADH conversion (i.e., in the absence of proteins) were used to compute the observed ATPase rates with the following formula:

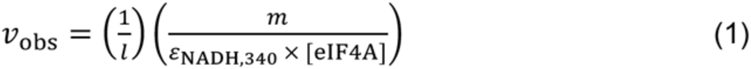

*l* is the pathlength, *m* is the background corrected slope, *ε*_NADH.340_ is the molar absorptivity of NADH at 340 nm, (6,220 mM^−1^ cm^−1^) and [eIF4A] is the total concentration of eIF4A added in the experiment. The reported values of *V* / *E*_o_ represent the average of three independent replicate experiments; error bars reflect the standard deviation of these replicates.

### Single-molecule experiments

A customized Pacific Biosciences RS II instrument was used to monitor real-time eIF4E interaction with the 5’ UTR of the viral RNA.^42,46,50^ For the experiment, a zero-mode waveguide (ZMW) chip (Pacific Biosciences SMRT Cell; part number 100-171-800) was first treated with NeutrAvidin mixture (16.6 µM neutravidin, 0.67 mg/mL BSA) in smBuffer 30 mM HEPES-KOH pH 7.5, 3 mM Mg(OAc)_2_ and 100 mM KOAc) at room temperature for 5 min, which allowed the NeutrAvidin to bind the biotin-PEG layer on base of the chip. Polyadenylated RNAs were prepared for immobilization through annealing to biotin-5’-(dT)_45_-3’-Cy3 (purchased from IDT), by heating a mixture containing 1 µM RNA and 100 nM oligo in 0.1 M BisTris pH 7.0, 0.3 M KCl to 98 °C for 2 minutes in a thermocycler, then slow-cooling to 4 °C over 20 minutes. The resulting hybrid duplex was immobilized by incubating 20 µL of the annealing mixture (10 nM RNA mixture final) on the chip surface for ∼10 min at room temperature. The immobilization mixture was then removed, and the chip was washed three times with smBuffer. The chip was then pre-blocked with 1 µM unlabeled eIF4E, 5% (v/v) each Biolipidure 203 and 206 (NOF America Corporation), and 5 mg/mL purified BSA (NEB, G9001S), to prevent non-specific interaction of Cy5-eIF4E with the chip surface. After pre-blocking, the chip was washed a further three times with smBuffer. After the third wash, 20 µL was added to the chip surface of smBuffer supplemented with BSA, with an oxygen scavenging system (2.5 mM PCA (protocatechuic acid), 1× PCD (protocatechuate-3,4-dioxygenase, Pacific Biosciences)), and with triplet-state quencher (2 mM TSY, Pacific Biosciences). After loading the chip onto the instrument, 15 nM Cy5 labeled hs4E WT was manually added to the chip, along with unlabeled components as needed (i.e., eIF4G, eIF4A, and ATP, ATP*γ*S), in a volume of 20 µL (final concentrations in the experiment are half the delivery concentrations, e.g. 7.5 nM eIF4E, due to dilution into the volume already on the chip surface). Where included, eIF4G(557-1137, 557-1600, 84-1600) and eIF4A were present at final concentrations of 25 nM and 1 µM, respectively. Movies were acquired for 10 minutes at 10 frames per second (fps) or 3 minutes at 80 fps, depending on the experiment, under illumination with a 532 nm laser at a power of 0.70 µW/µm^2^, to excite Cy3 fluorophores. In experiments with direct red illumination, the 642 nm laser power was used at a power of 0.07 µW/µm^2^.

### Single-molecule data analysis

Single-molecule fluorescence traces were extracted from raw movie files with custom MATLAB scripts reported previously.^46,58^ Traces containing robust smFRET signals for eIF4E–RNA interaction were selected by visual inspection. Selection criteria were: a stable Cy3 signal at the very beginning of the movie, a single Cy3 photobleaching event, and visually apparent FRET to Cy5. Assignment of event timings in the traces was carried out using a hidden Markov model approach. Event timings were converted to empirical cumulative probability distributions for the times between events, and the event durations, then fit with a single-exponential model to determine *k*_on_ or *k*_off_, according to the equation:

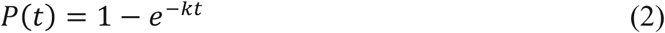

Fitting was carried out in MATLAB by non-linear least-squares regression using the Trust-Region algorithm. Goodness-of-fit was confirmed by inspection of the root-mean-squared errors for the fits, which were 0.01 – 0.03 in all cases.

### Molecular dynamics simulations

Molecular dynamics simulations were performed using the HOOMD-blue package.^76^ Proteins were modeled as rigid bodies, while RNA molecules were represented using a spring-bead model incorporating secondary structures predicted by the ViennaRNA package. A poly(A) tail, 45a in length (with 1a corresponding to 1nm), was appended to the RNA’s 3’ end. At the 5’ end, a cap structure was modeled as a distinct particle. The bead diameters for mRNA, poly(A), and cap particles were set to 0.33a, 0.33a, and 0.5a, respectively, with corresponding charges of −0.5, −0.5, and −1.5. Protein-RNA interactions were modeled by the Debye–Hückel potential, which can be written as,

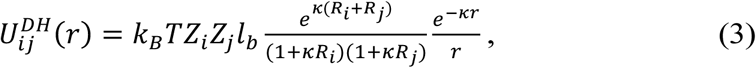

where the Debye screening length is *k*^−1^ = 1*nm*. The Bjerrum length, defined as 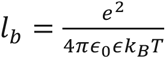 quantifies the distance at which the electrostatic interaction between two elementary charges equals the thermal energy *k*_*B*_*T*. For water at room temperature, *l*_*b*_ ≈ 0.7*nm*. The interaction between the protein and the cap is modeled using the Lennard-Jones (LJ) potential, which can be written as,

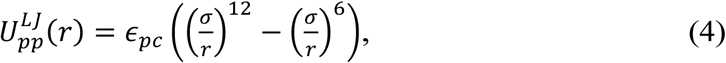

where *ϵ*_*pc*_ = 8*k*_*B*_*T* represents the interaction strength, *σ* = 2^−1/6^(*R*_*i*_ + *R*_j_) ensures that the potential minimum occurs at the sum of the particle radii, with *R*_*i*_ and *R*_j_ being the radii of particles i and j, respectively. r denotes the center-to-center distance between the interacting particles. The interaction is truncated at a cutoff distance of *r*_*cut*_ = 1.6*a*, where the unit length *a* is defined as 1*nm*. All other particles in the system interact through the repulsive Lennard-Jones potential,

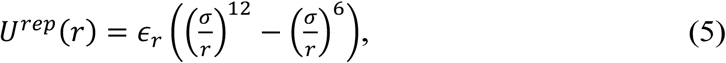

which is subject to a cutoff distance *r*_*cut*_ = *R*_*i*_ + *R*_j_, with an interaction strength of *ϵ*_*r*_ = 0.1*k*_*B*_*T*.

Protein dynamics are simulated using a Langevin integrator with a time step of *dt* = 0.0005*τ*, where *τ* is the fundamental time unit of the system, approximately equivalently to 1 second. Simulation snapshots are recorded every 2000 steps. For each type of protein and RNA, five independent simulation runs are performed to obtain ensemble data.

In the simulation, binding events are detected by monitoring the distance between the center of the protein and the cap particle. If the distance falls below 5 nm, the state is marked as ’1’ (bound); otherwise, it is marked as ’0’ (unbound). This generates a binary time series—an on/off barcode— over the course of 5,000 simulation frames.

To quantify binding durations, we extract all ’on’ intervals by identifying transitions from ’0’ to ’1’ (binding onset) and from ’1’ to ’0’ (binding termination). The duration of each event is calculated by subtracting the start time from the end time. These event durations are compiled to construct a cumulative distribution function (CDF), which is then fit to a double exponential model.

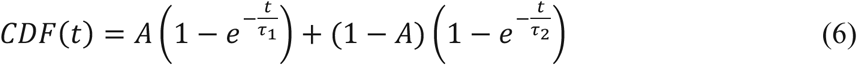

where *τ*_1_ and *τ*_2_ are characteristic time constants, and *A* is a weighting factor. We note that for the association rate, we use *τ*_1_, and for the dissociation rate, we use *τ*_2_as these values are more consistent with the experimental data.

## ACKNOWEDLGEMENTS

Plasmid pULTRA for unnatural amino acid incorporation was a gift from Abhishek Chatterjee (Boston College). Plasmid pHis-APEX2-eIF4A1 was a gift from Nicholas Ingolia (University of California Berkeley; Addgene plasmid #129645).

## AUTHOR CONTRIBUTIONS

This work was conceptualised by S.O’L, H.H. and R.Z.. Single-molecule experiments and biochemical analyses were performed, analysed and interpreted by H.H., MG., A.H. and E.L. with input from S.O’L. R.H. designed the plasmid construct for human eIF4G expression and A.N. and D.X. generated the resulting human cell line with input from R.H. S.L. performed, analysed and interpreted the molecular dynamics simulations with input from R.Z. and S.O’L. Y.L. developed the analytical calculation with input from R.Z. H.H., SO’L and R.Z. wrote the paper.

## DATA AVAILABILITY

All data analysed in this study are available in the manuscript and Extended Data.

## CODE AVAILABILITY

MATLAB code for single-molecule data analysis is available at: https://drive.google.com/drive/folders/1k1ZvZqb-TMgjpysfUVnEEwbiFJ7s0rsr?usp=share_link

---

**Extended Data Figure 1.**
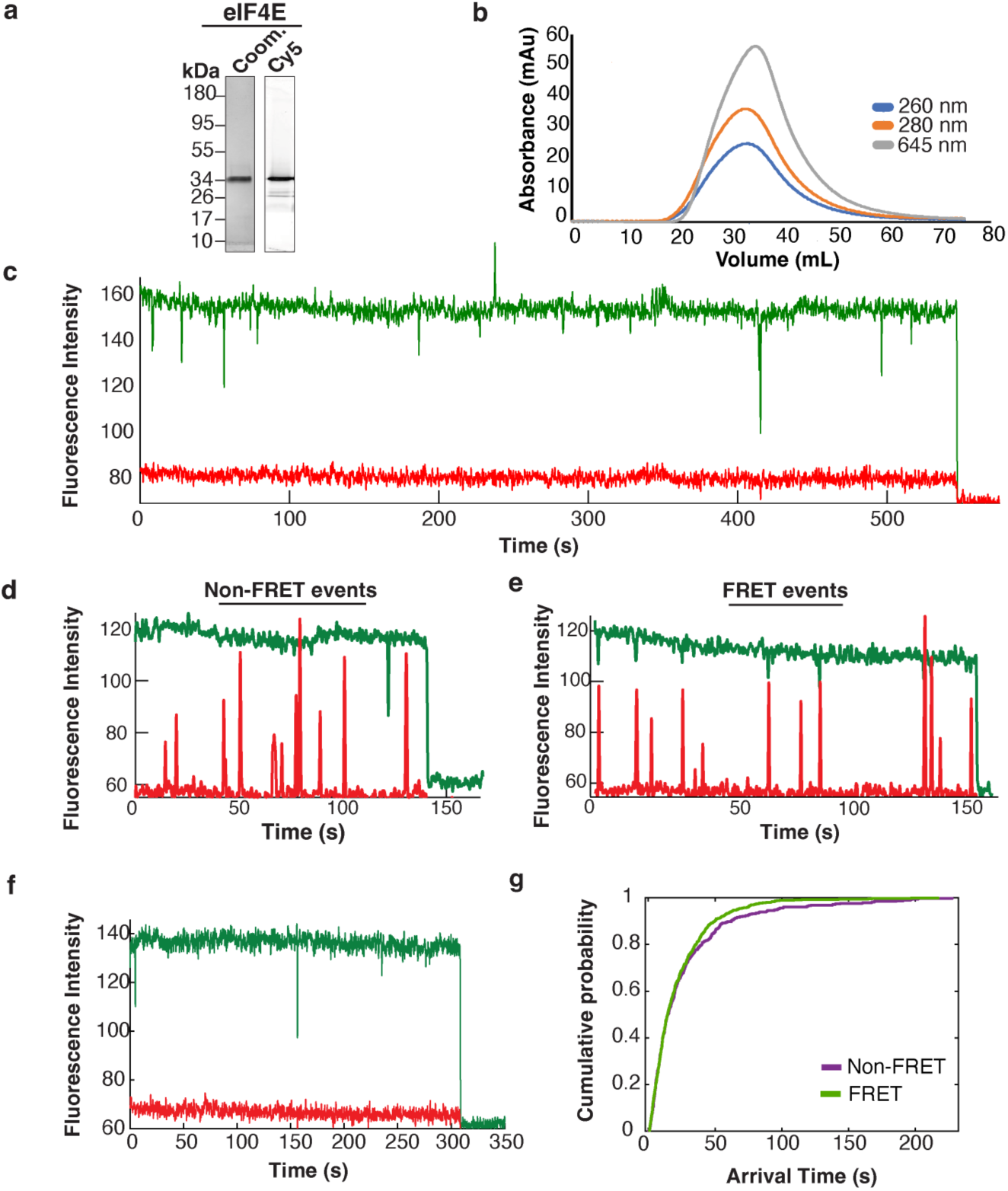
Purification and validation of human eIF4E a. SDS-PAGE analysis of purified eIF4E. The gel on the left is stained with Coomassie blue, and the right is a Typhoon image taken prior to Coomassie staining. **b.** Size-exclusion chromatogram from eIF4E purification on Superdex 75. **c.** Representative trace for eIF4E interaction with the viral RNA in the presence of 1 mM m^7^GpppG cap analog. **d.** Representative trace for non-FRET eIF4E interaction with the viral RNA, under continuous illumination with both green and red lasers, which directly detects Cy5-eIF4E binding. **e.** Representative trace from an RNA molecule where eIF4E interaction dominantly results in FRET, even though all interactions are detected with simultaneous green/red illumination. **f.** Representative trace of eIF4E interaction with a *β*-globin mRNA in the presence of 1 mM m^7^GpppG and with dual red/green illumination. **g.** Cumulative distribution functions for eIF4E–cap association for FRET (green) and non-FRET (purple) events.

**Extended Data Figure 2.**
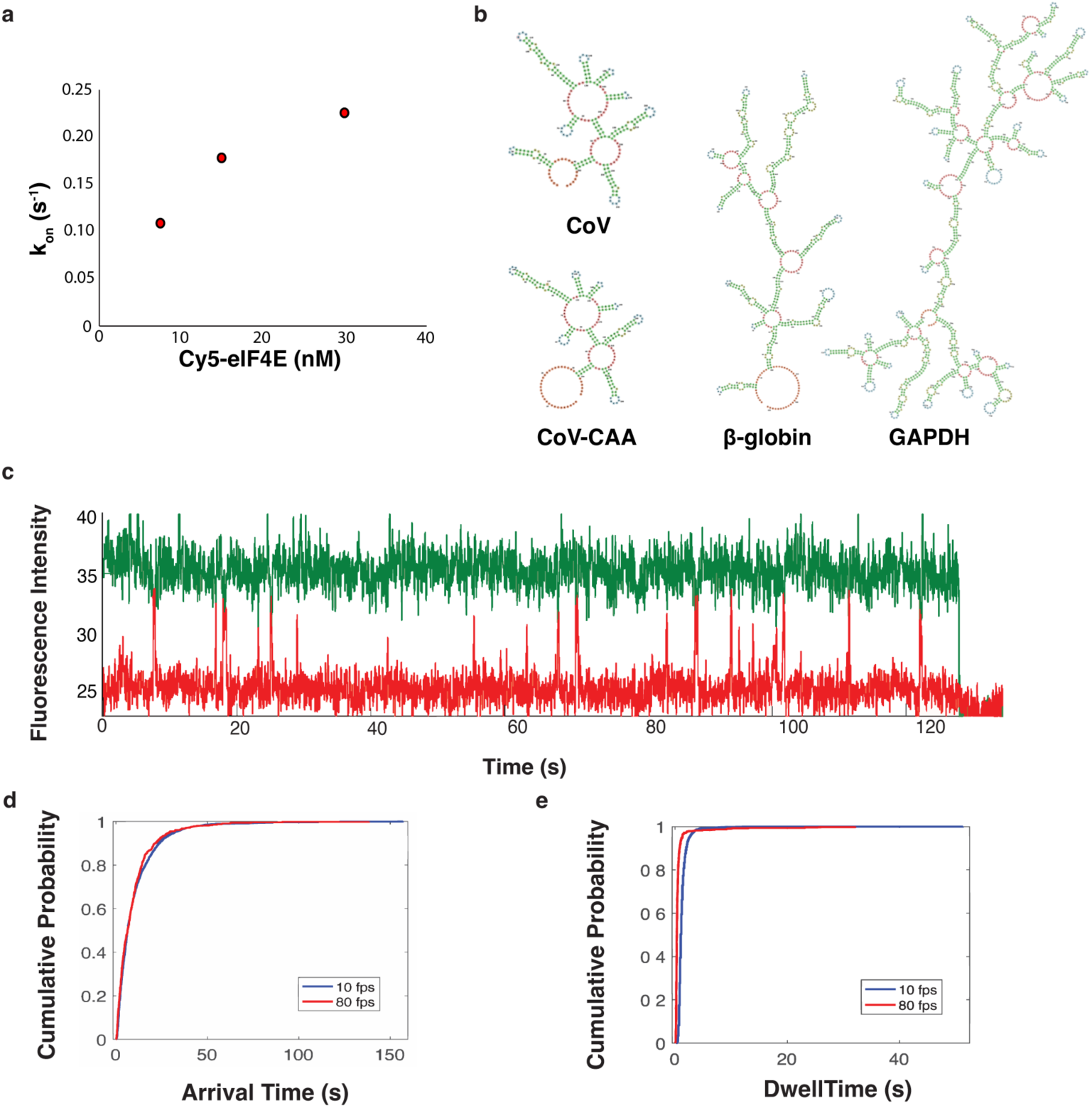
**a.** Dependence of human Cy5-eIF4E–CoV RNA association rate on Cy5-eIF4E concentration. **b.** Computational folding prediction of CoV, CoV-CAA, *β*-globin, GAPDH RNAs. **c.** Representative trace for Cy5-eIF4E interaction with the viral RNA, with data acquisition at 80 fps. **d-e.** Cumulative distribution functions for measured eIF4E–cap arrival (**d**) and dwell times (**e**) at 10 (blue) and 80 fps (red).

**Extended Data Figure 3.**
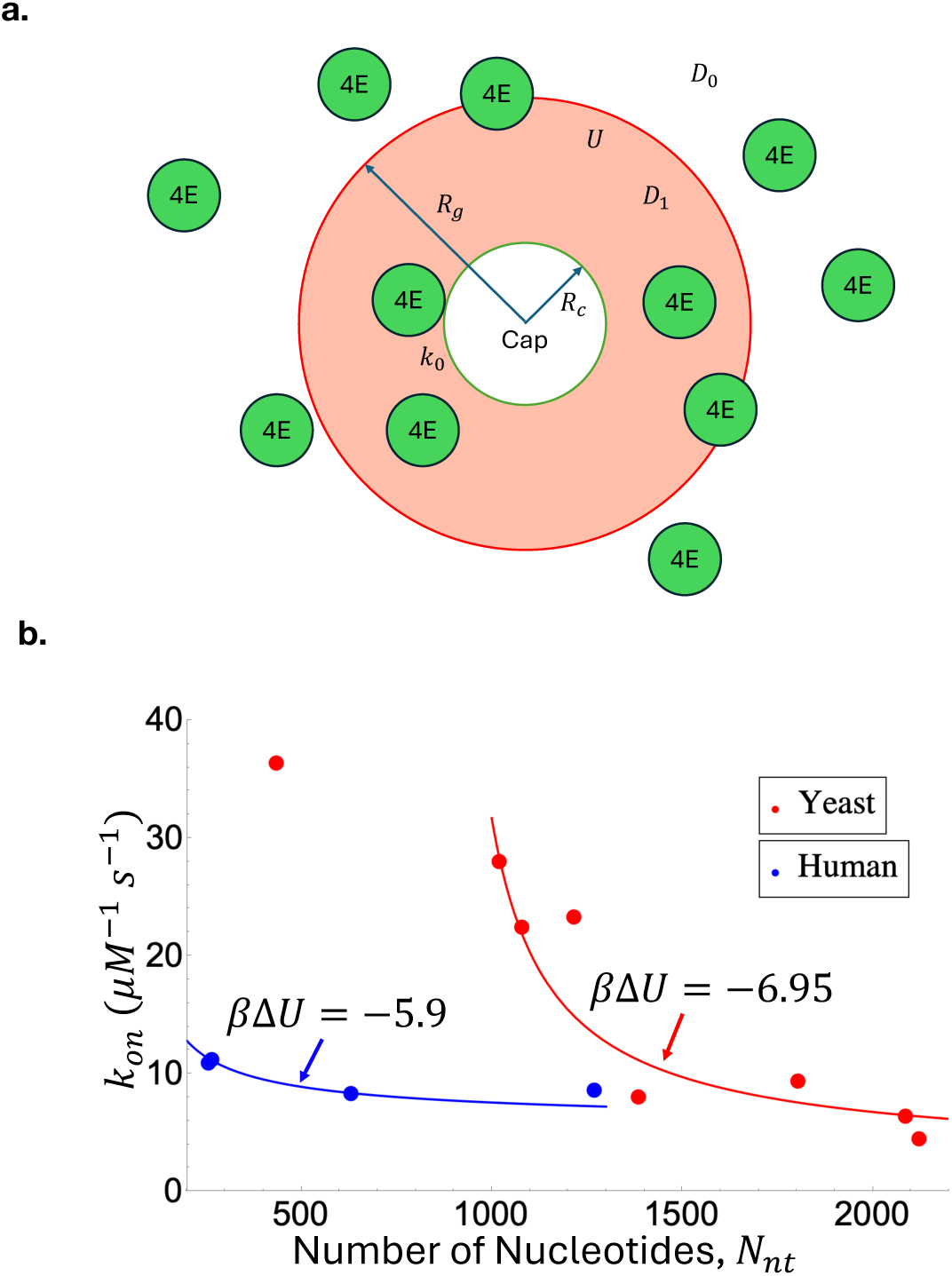
**a.** Schematic plot of the diffusion-reaction-based model. The RNA cap is modeled as a sphere of radius *R*_*c*_ and binds to eIF4E with an intrinsic/reaction-controlled rate constant *k*_0_when they come within the reaction range. The radius of gyration of the RNA chain is denoted as *R*_g_. The diffusion coefficient of eIF4E is *D*_0_ when it is outside *R*_g_and *D*_1_when it is within *R*_g_. *U* is the effective potential that eIF4E experiences within *R*_g_. *k*_*on*_ vs *N*_*nt*_ for human (blue) and yeast (red) eIF4E. **b.** The fitting yields *D*_0_ = 0.71*μm*^2^/*s*, *D*_1_ = 184µ*m*^2^/*s*, *R*_*c*_ = 1.26*nm* and βΔ*U* = −6.95.

**Extended Data Figure 4.**
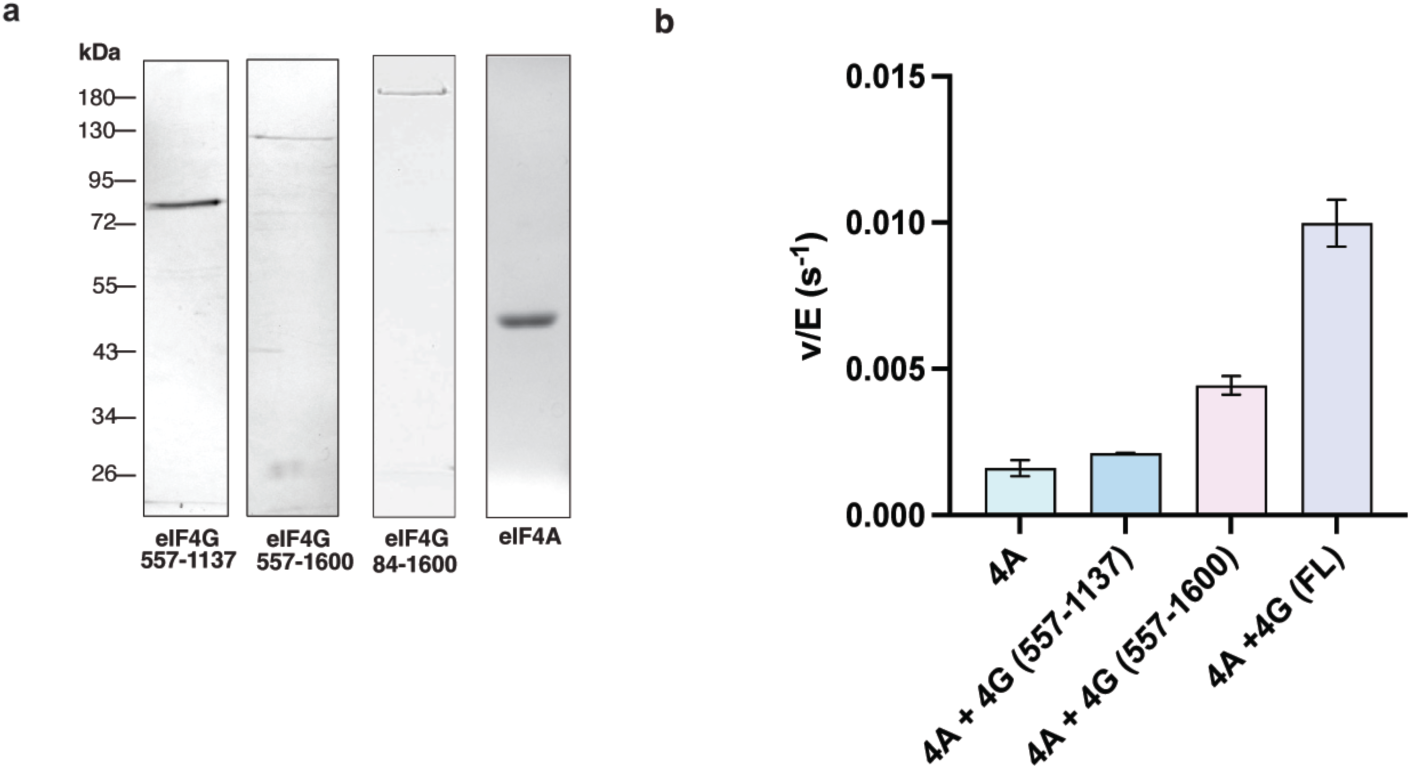
**a.** SDS-PAGE analysis of purified human eIF4G proteins (557-1137, 557-1600, 84-1600). **b.** Stimulation of eIF4A (500 nM) ATPase activity by eIF4G (557-1137, 12.5 nM; 557-1600, 12.5 nM; FL or 84-1600, 12.5 nM) determined by an NADH-linked coupled spectrophotometric assay.

**Extended Data Figure 5.**
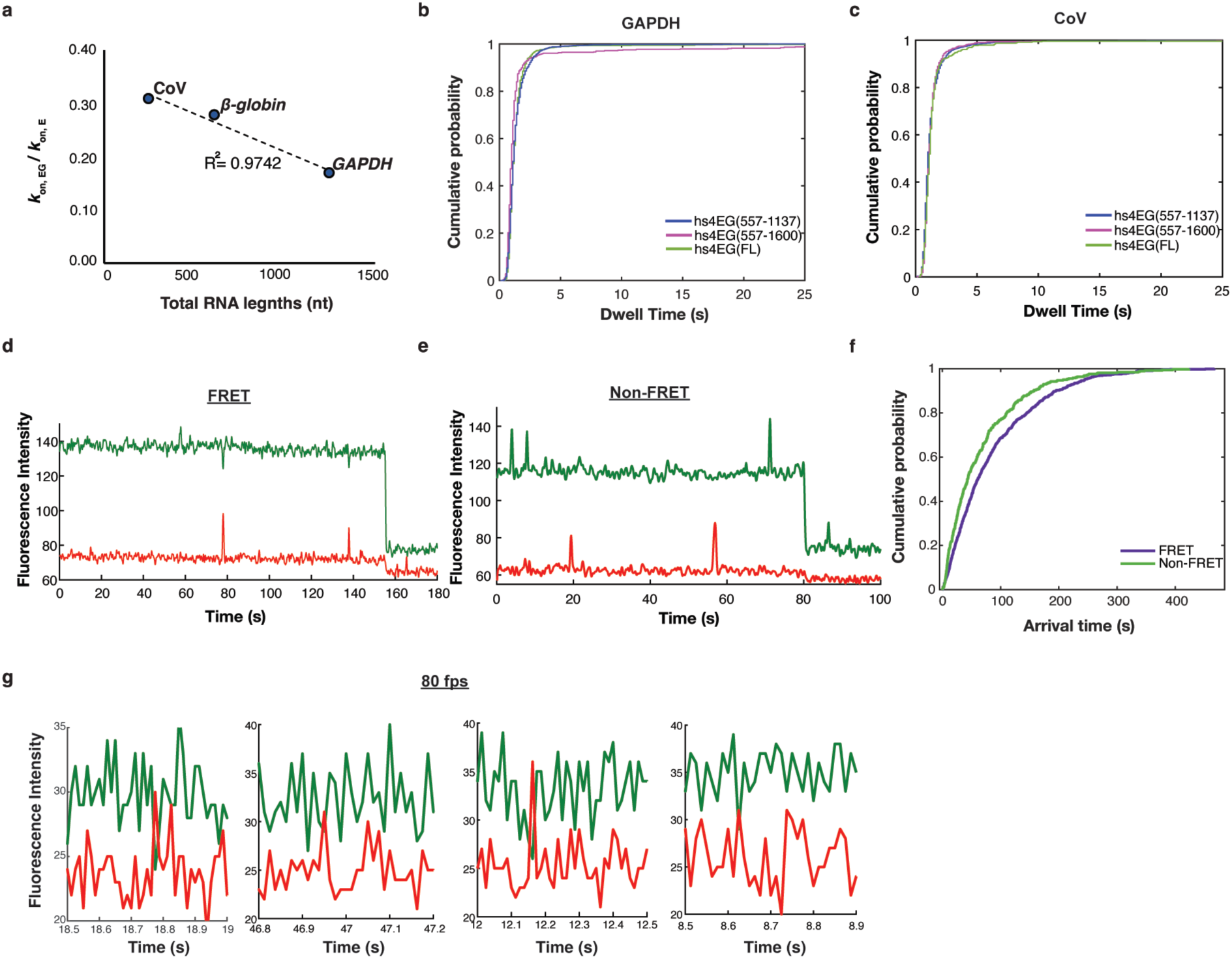
**a.** Fold change in eIF4E–mRNA association rate resulting from addition of eIF4G, plotted as a function of mRNA total length. **b.** Cumulative distribution functions for eIF4E–cap dwell times in the presence of eIF4G(557-1137/557-1600/84-1600) on *GAPDH* mRNA. **c.** Cumulative distribution functions for eIF4E–cap dwell time in the presence of eIF4G(557-1137/557-1600/84-1600) on CoV RNA. **d-e.** Representative traces of eIF4E•G interaction (**d.** FRET or **e.** FRETless) with the viral RNA, under conditions of dual red-green laser illumination. **f.** Cumulative distribution functions for eIF4E– cap association in the presence of eIF4G(84-1600) on CoV RNA for FRET (green) and non-FRET (purple) events. **g.** Representative traces for eIF4E•G interaction with the *GAPDH* mRNA at 80 fps, under conditions of dual red-green laser illumination.

**Extended Data Figure 6.**
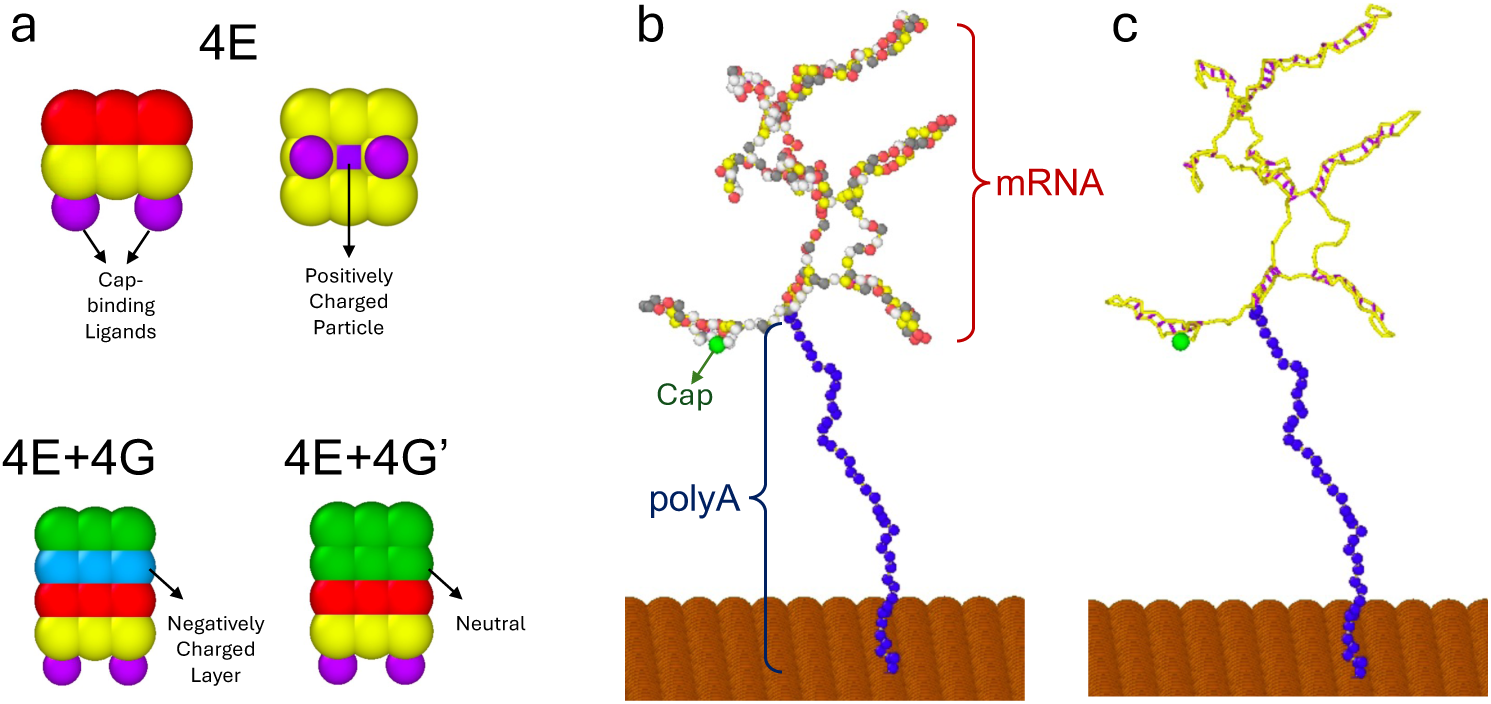
**a.** Schematic representations of eIF4E, eIF4E–eIF4G, and eIF4E–eIF4G′ complexes. eIF4E is depicted as two layers of beads: red and yellow. eIF4G is shown with a negatively charged blue layer and a neutral green layer. eIF4G′, which mimics the role of eIF4A by neutralizing some of the charges in eIF4G, consists of two neutral green layers. The purple beads have a radius of 0.35a, while all other beads have a radius of 0.5a. The positively charged particle in eIF4E carries a charge of +5, whereas each negatively charged bead in eIF4E-eIF4G carries a charge of -1. **b.** Diagram illustrating mRNA molecules anchored on a rigid surface. A 45a-length poly(A) tail (blue) is appended to the 3’ end of RNA. A green cap particle is attached at the 5’ end. The bead diameters for mRNA, poly(A) tail, and cap particles are 0.33a, 0.33a, and 0.5a, respectively. The corresponding charges are −0.5, −0.5, and −1.5. **c.** Visualization of intra-molecular mRNA base-pairing interactions representing secondary structure formation.

## Supplementary Note

Analytical Calculation

The kinetics of the single-molecule reaction are modeled through a diffusion-reaction-based model, as illustrated in Extended Data Figure 3a.

The RNA cap is assumed to be spherical with radius *R*_*c*_. The intrinsic/reaction-controlled rate constant *k*_0_ governs the cap-eIF4E binding process when they come within the reaction range. We assume *k*_0_ is sufficiently large such that the time scale of this reaction is much smaller than the time resolution in the experiment, effectively making the binding instantaneous upon touch. The RNA chain has a radius of gyration *R*_g_. According to Flory scaling law, we have 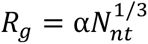, where α is the length scale of RNA in unit nm, and *N*_*nt*_ is the number of nucleotides. Within the radius of gyration of the RNA, the diffusion coefficient of eIF4E is *D*_1_, which is different from the diffusion coefficient *D*_0_ when eIF4E is away from the RNA. The difference comes from the interactions between eIF4E and both the RNA chain and the RNA cap, such as the excluded volume effect and electrostatic interactions. These interactions not only modify the diffusion directly by altering the diffusion coefficient, but also generate a potential of mean force, which we assume to be uniform and denote by *U*. For example, the attractive force between eIF4E and the RNA cap will create a negative potential field, as one contribution to *U*. Note that we have omitted the varying interaction ranges for different interactions for simplicity. In this model, such interactions only happen within *R*_g_, thus *U* = 0 when outside of *R*_g_.

Following Ref. [1], the association rate between eIF4E and the RNA cap *k*_*on*_ is expressed as

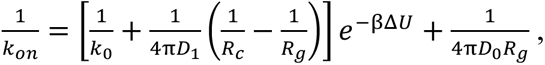

where β = (*k*_*B*_*T*)^−1^, *k*_*B*_ is the Boltzmann’s constant, *T* is the absolute temperature and Δ*U* = *U* − 0. In the experiments, the association rate *k*_*on*_is measured for different RNA chains with varying lengths, suggesting us to treat *k*_*on*_as a function of *R*_g_ and fit the other parameters. To determine the parameters, we first apply the analytical model to our previous data for yeast RNAs in Ref. [2]. The corresponding results are presented in red in Extended Data Figure 3b.

The fitting yields *D*_0_ = 0.71*μm*^2^/*s*, *D*_1_ = 184µ*m*^2^/*s*, *R*_*c*_ = 1.26*nm* and βΔ*U* = −6.95 . We have taken *k*_0_ to be sufficiently large following our previous assumption and α = 0.1*nm* since the fitting is not sensitive to this parameter. The comparison between the two diffusion coefficients indicates that the mobility of eIF4E is significantly enhanced near the cap, despite the potential hindrance from the RNA body. The negative potential indicates the depletion effect, leading to the decreased local concentration of eIF4E within the range *R*_g_, as discussed in Ref. [3]. Our model successfully captures the length dependence of the association rate for longer segments of yeast RNAs. For human RNAs, which are the focus of the current paper, the length dependence is weaker. Notably, our model still fits the data well by adjusting only one parameter: *β*Δ*U* = −5.9, with *D*_0_, *D*_1_ and *R*_*c*_ unchanged. The reduced magnitude of the potential suggests a weaker depletion interaction for human RNAs.

Overall, our model reproduces key trends of the association rate as a function of RNA length, and may provide insight into the mechanism in the experiment.

## Supplementary Information

DNA construct sequences

All coding sequences for bacterial expression are codon-optimized. Sequences are inserted into plasmid sites as indicated in the methods section. The following sequences are presented in a 5’ to 3’ direction, with initiation and the stop codons underlined. DNA template sequences for RNA constructs are also shown in the 5’ to 3’ direction. Restriction sites are indicated in bold, and the T7 promoter sequence is underlined.

### SARS-CoV-2 UTR

**GGATCC**<u>TAATACGACTCACTATT</u>ATTAAAGGTTTATACCTTCCCAGGTAACAAA CCAACCAACTTTCGATCTCTTGTAGATCTGTTCTCTAAACGAACTTTAAAATCT GTGTGGCTGTCACTCGGCTGCATGCTTAGTGCACTCACGCAGTATAATTAATAA CTAATTACTGTCGTTGACAGGACACGAGTAACTCGTCTATCTTCTGCAGGCTGC TTACGGTTTCGTCCGTGTTGCAGCCGATCATCAGCACATCTAGGTTTCGTCCGG GTGTGACCGAAAGGTAAG**GAATTC**

### SARS-CoV-2 CoV-CAA

**GGATCC**<u>TAATACGACTCACTATT</u>ATTAAACAACAACAACAACAACAAAACCAA CTTTCGATCTCTTGTAGATCTGTTCTCTAAACGAACTTTAAAATCTGTGTGGCTGTCACTCGGCTGCATGCTTAGTGCACTCACGCAGTATAATTAATAACTAATTACTGTCGTTGACAGGACACGAGTAACTCGTCTATCTTCTGCAGGCTGCTTACGGTTTCGTCCGTGTTGCAGCCGATCATCAGCACATCTAGGTTTCGTCCGGGTGTGAC CGAAAGGTAAG**GAATTC**

#### GAPDH

**GTCGAC**<u>TAATACGACTCACTATAG</u>GTGTTCGACAGTCAGCCGCATCTTCTTTTG CGTCGCCAGCCGAGCCACATCGCTCAGACACCATGGGGAAGGTGAAGGTCGGAGTCAACGGATTTGGTCGTATTGGGCGCCTGGTCACCAGGGCTGCTTTTAACTCTGGTAAAGTGGATATTGTTGCCATCAATGACCCCTTCATTGACCTCAACTACATGGTTTACATGTTCCAATATGATTCCACCCATGGCAAATTCCATGGCACCGTCAAGGCTGAGAACGGGAAGCTTGTCATCAATGGAAATCCCATCACCATCTTCCAGGAGTGAGTGGAAGACAGAATGGAAGAAATGCGAGATCCCTCCAAAATCAAGTGGGGCGATGCTGGCGCTGAGTACGTCGTGGAGTCCACTGGCGTCTTCACCACCATGGAGAAGGCTGGGGCTCATTTGCAGGGGGGAGCCAAAAGGGTCATCATCTCTGCCCCCTCTGCTGATGCCCCCATGTTCGTCATGGGTGTGAACCATGAGAAGTATGACAACAGCCTCAAGATCATCAGCAATGCCTCCTGCACCACCAACTGCTTAGCACCCCTGGCCAAGGTCATCCATGACAACTTTGGTATCGTGGAAGGACTCATGACCACAGTCCATGCCATCACTGCCACCCAGAAGACTGTGGATGGCCCCTCCGGGAAACTGTGGCGTGATGGCCGCGGGGCTCTCCAGAACATCATCCCTGCCTCTACTGGCGCTGCCAAGGCTGTGGGCAAGGTCATCCCTGAGCTGAACGGGAAGCTCACTGGCATGGCCTTCCGTGTCCCCACTGCCAACGTGTCAGTGGTGGACCT GACCTGCCGTCTAGAAAAACCTGCCAAATATGATGACATCAAGAAGGTGGTGAAGCAGGCGTCGGAGGGCCCCCTCAAGGGCATCCTGGGCTACACTGAGCACCAGGTGGTCTCCTCTGACTTCAACAGCGACACCCACTCCTCCACCTTTGACGCTGGGGCTGGCATTGCCCTCAACGACCACTTTGTCAAGCTCATTTCCTGGTATGACAACGAATTTGGCTACAGCAACAGGGTGGTGGACCTCATGGCCCACATGGCCTCCAAGGAGTAAGACCCCTGGACCACCAGCCCCAGCAAGAGCACAAGAGGAAGAGAGAGACCCTCACTGCTGGGGAGTCCCTGCCACACTCAGTCCCCCACCACACTGAATCTCCCCTCCTCACAGTTGCCATGTAGACCCCTTGAAGAGGGGAGGGG CCTAGGGAGCCGCACCTTGTCATG**GAATTC**

#### β-Globin

**AAGCTT**<u>TAATACGACTCACTATAG</u>GACATTTGCTTCTGACACAACTGTGTTCAC TAGCAACCTCAAACAGACACCATGGTGCATCTGACTCCTGAGGAGAAGTCTGCCGTTACTGCCCTGTGGGGCAAGGTGAACGTGGATGAAGTTGGTGGTGAGGCCCTGGGCAGGCTGCTGGTGGTCTACCCTTGGACCCAGAGGTTCTTTGAGTCCTTTGGGGATCTGTCCACTCCTGATGCTGTTATGGGCAACCCTAAGGTGAAGGCTCATGGCAAGAAAGTGCTCGGTGCCTTTAGTGATGGCCTGGCTCACCTGGACAACCTCAAGGGCACCTTTGCCACACTGAGTGAGCTGCACTGTGACAAGCTGCACGTGGATCCTGAGAACTTCAGGCTCCTGGGCAACGTGCTGGTCTGTGTGCTGGCCCATCACTTTGGCAAAGAATTCACCCCACCAGTGCAGGCTGCCTATCAGAAAGTGGTGGCTGGTGTGGCTAATGCCCTGGCCCACAAGTATCACTAAGCTCGCTTTCTTGCTGTCCAATTTCTATTAAAGGTTCCTTTGTTCCCTAAGTCCAACTACTAAACTGGGGGATATTATGAAGGGCCTTGAGCATCTGGATTCTGCCTAATAAAAAACA TTTATTTTCATTGCAA**GTCGAC**

### Human eIF4E

**CC<u>ATG</u>G**GTAGCTCACATCATCATCATCATCACTCTTCTGGTCTGGTCCCGCGTG GCTCGCACATGCAATACAAACTGATTCTGAACGGTAAAACGCTGAAAGGTGAAACCACGACCGAAGCAGTGGATGCGGCCACCGCTGAAAAAGTTTTCAAACAGTACGCCAACGATAATGGCGTGGATGGTGAATGGACCTATGATGACGCAACGAAAACCTACACGGTGACCGAAGGTTCCGGCGGTGAAAATCTGTACTTCCAAGGCCATATGGCCACCGTTGAACCGGAAACGACCCCGACGACCAACCCGCCGCCGGCTGAAGAAGAAAAAACCGAAAGCAACCAGGAAGTCGCGAATCCGGAACATTATATTAAACACCCGCTGCAAAACCGTTGGGCTCTGTGGTTTTTCAAAAACGATAAATCAAAAACGTGGCAGGCGAACCTGCGCCTGATTTCGAAATTTGATACCGTGGAAGACTTCTGGGCACTGTATAACCACATCCAACTGAGCTCTAATCTGATGCCGGGTTGCGATTACAGCCTGTTTAAAGACGGCATTGAACCGATGTGGGAAGATGAGAAAAACAAACGTGGCGGTCGCTGGCTGATCACGCTGAACAAACAGCAACGTCGCTCTGATCTGGACCGTTTTTGGCTGGAAACCCTGCTGTGCCTGATTGGCGAAAGTTTCGATGACTACTCCGATGACGTTTGTGGTGCGGTGGTTAATGTCC GTGCCAAAGGCGATAAAATTGCAATCTGGACGACCGAATGTGAAAACCGCGACGCCGTCACCCATATCGGCCGTGTGTATAAAGAACGCCTGGGTCTGCCGCCGAAAATTGTTATCGGCTACCAGAGCCACGCAGATACGGCGACCAAATCGGGCAGC ACCACCAAAAATCGTTTCGTTGTG<u>TGA</u>**CTCGAG**

### Human eIF4G (557-1137)

**CC<u>ATG</u>G**GGCATCATCACCATCACCACGAGTCAGAAGGGTCGGGAGTCCCGCC AAGACCCGAAGAGGCGGATGAGACATGGGATTCGAAAGAGGATAAAATCCATAATGCTGAAAATATACAACCTGGAGAGCAGAAATACGAGTATAAATCGGACCAATGGAAACCATTGAACTTAGAGGAAAAAAAGCGCTACGATCGTGAATTCCTTTTGGGGTTTCAATTCATCTTTGCATCGATGCAAAAGCCAGAGGGCCTGCCACATATCAGTGATGTTGTACTTGACAAGGCGAATAAGACACCGTTAAGACCCTTAGATCCTACGCGTTTACAAGGAATAAATTGCGGACCAGATTTTACGCCTAGCTTCGCAAATCTTGGTAGAACAACCCTGAGCACACGCGGCCCTCCCAGAGGTGGCCCAGGGGGGGAGCTGCCACGCGGACCAGCAGGTTTAGGGCCAAGACGTTCACAACAGGGGCCGCGGAAAGAACCTCGCAAAATCATAGCGACGGTATTGATGACAGAGGATATTAAATTGAATAAAGCAGAAAAGGCGTGGAAGCCATCTTCGAAACGCACAGCCGCTGATAAGGATCGGGGTGAGGAGGACGCCGATGGCAGCAAAACACAAGATCTTTTCCGGCGGGTCCGTAGCATATTAAACAAGCTTACTCCACAGATGTTCCAACAGCTGATGAAGCAAGTGACACAGTTGGCGATTGACACAGAAGAACGGCTTAAGGGGGTCATTGATTTGATATTTGAAAAAGCCATTAGTGAGCCCAACTTCAGCGTAGCTTACGCTAATATGTGTCGTTGTTTAATGGCGCTTAAAGTTCCGACTACGGAGAAACCGACCGTAACAGTGAATTTCCGTAAATTGCTTTTAAACCGTTGTCAAAAGGAATTCGAAAAAGACAAGGATGACGACGAGGTTTTCGAGAAAAAACAAAAAGAGATGGACGAGGCCGCTACAGCAGAGGAAAGAGGTAGATTAAAGGAAGAATTAGAAGAGGCCCGCGACATCGCGCGGCGGAGAAGCTTGGGAAATATCAAATTTATAGGGGAACTTTTCAAATTGAAAATGCTGACTGAGGCAATCATGCATGACTGCGTTGTTAAGCTGTTAAAGAACCATGACGAGGAGTCGTTAGAGTGCTTGTGCCGGTTACTTACAACAATTGGCAAAGATCTGGACTTTGAAAAGGCGAAGCCACGCATGGACCAATACTTTAACCAGATGGAAAAAATTATAAAGGAAAAAAAAACCTCGAGCCGGATAAGATTTATGTTGCAAGACGTACTGGATTTACGTGGGTCTAACTGGGTTCCCCGTCGGGGGGACCAGGGTCCGAAGACCATAGATCAAATCCACAAGGAAGCGGAAATGGAAGAGCATCGTGAACATATAAAGGTTCAACAATTAATGGCCAAAGGGTCAGACAAGCGGCGTGGCGGTCCCCCTGGGCCGCCCATCTCAAGAGGATTACCATTGGTTGACGACGGCGGATGGAATACCGTACCTATTAGCAAGGGCAGCCGCCCTATAGACACATCGCGGCTTACTAAGATTACGAAACCAGGGTCGATTGATTCCAACAACCAACTGTTTGCACCAGGCGGTCGCTTGAGTTGGGGCAAGGGTTCATCGGGCGGTTCGGGAGCAAAGCCCTCTGATGCCGCCTCGGAAGCAGCGAGACCAGCAACGTCTACTCTTAACCGGTTTTCTGCTCTGC AACAAGCGGTCCCTACCGAGTCTACGGATAAC<u>TA**G**</u>**GATCC**

### Human eIF4G (557-1600)

**CC<u>ATG</u>G**GCCATCATCATCATCATCATGAGTCTGAGGGCAGTGGTGTGCCCCCA CGTCCTGAGGAAGCAGATGAGACCTGGGACTCAAAGGAAGACAAAATTCACAATGCTGAGAACATCCAGCCCGGGGAACAGAAGTATGAATATAAGTCAGATCAGTGGAAGCCTCTAAACCTAGAGGAGAAAAAACGTTACGACCGTGAGTTCCTGCTTGGTTTTCAGTTCATCTTTGCCAGTATGCAGAAGCCAGAGGGATTGCCACATATCAGTGACGTGGTGCTGGACAAGGCCAATAAAACACCACTGCGGCCACTGGATCCCACTAGACTACAAGGCATAAATTGTGGCCCAGACTTCACTCCATCCTTTGCCAACCTTGGCCGGACAACCCTTAGCACCCGTGGGCCCCCAAGGGGTGGGCCAGGTGGGGAGCTGCCCCGTGGGCCGGCTGGCCTGGGACCCCGGCGCTCTCAGCAGGGACCCCGAAAAGAACCACGCAAGATCATTGCCACAGTGTTAATGACCGAAGATATAAAACTGAACAAAGCAGAGAAAGCCTGGAAACCCAGCAGCAAGCGGACGGCGGCTGATAAGGATCGAGGGGAAGAAGATGCTGATGGCAGCAAAACCCAGGACCTATTCCGCAGGGTGCGCTCCATCCTGAATAAACTGACACCCCAGATGTTCCAGCAGCTGATGAAGCAAGTGACGCAGCTGGCCATCGACACCGAGGAACGCCTCAAAGGGGTCATTGACCTCATTTTTGAGAAGGCCATTTCAGAGCCCAACTTCTCTGTGGCCTATGCCAACATGTGCCGCTGCCTCATGGCGCTGAAAGTGCCCACTACGGAAAAGCCAACAGTGACTGTGAACTTCCGAAAGCTGTTGTTGAATCGATGTCAGAAGGAGTTTGAGAAAGACAAAGATGATGATGAGGTTTTTGAGAAGAAGCAAAAAGAGATGGATGAAGCTGCTACGGCAGAGGAACGAGGACGCCTGAAGGAAGAGCTGGAAGAGGCTCGGGACATAGCCCGGCGGCGCTCTTTAGGGAATATCAAGTTTATTGGAGAGTTGTTCAAACTGAAGATGTTAACAGAGGCAATAATGCATGACTGTGTGGTCAAACTGCTTAAGAACCATGATGAAGAGTCCCTTGAGTGCCTTTGTCGTCTGCTCACCACCATTGGCAAAGACCTGGACTTTGAAAAAGCCAAGCCCCGAATGGATCAGTATTTCAACCAGATGGAAAAAATCATTAAAGAAAAGAAGACGTCATCCCGCATCCGCTTTATGCTGCAGGACGTGCTGGATCTGCGAGGGAGCAATTGGGTGCCACGCCGAGGGGATCAGGGTCCCAAGACCATTGACCAGATCCATAAGGAGGCTGAGATGGAAGAACATCGAGAGCACATCAAAGTGCAGCAGCTCATGGCCAAGGGCAGTGACAAGCGTCGGGGCGGTCCTCCAGGCCCTCCCATCAGCCGTGGACTTCCCCTTGTGGATGATGGTGGCTGGAACACAGTTCCCATCAGCAAAGGTAGCCGCCCCATTGACACCTCACGACTCACCAAGATCACCAAGCCTGGCTCCATCGATTCTAACAACCAGCTCTTTGCACCTGGAGGGCGACTGAGCTGGGGCAAGGGCAGCAGCGGAGGCTCAGGAGCCAAGCCCTCAGACGCAGCATCAGAAGCTGCTCGCCCAGCTACTAGTACTTTGAATCGCTTCTCAGCCCTTCAACAAGCGGTACCCACAGAAAGCACAGATAATAGACGTGTGGTGCAGAGGAGTAGCTTGAGCCGAGAACGAGGCGAGAAAGCTGGAGACCGAGGAGACCGCCTAGAGCGGAGTGAACGGGGAGGGGACCGTGGGGACCGGCTTGATCGT GCGCGGACACCAGCTACCAAGCGGAGCTTCAGCAAGGAAGTGGAGGAGCGGAGTAGAGAACGGCCCTCCCAGCCTGAGGGGCTGCGCAAGGCAGCTAGCCTCACGGAGGATCGGGACCGTGGGCGGGATGCCGTGAAGCGAGAAGCTGCCCTACCCCCAGTGAGCCCCCTGAAGGCGGCTCTCTCTGAGGAGGAGTTAGAGAAGAAATCCAAGGCTATCATTGAGGAATATCTCCATCTCAATGACATGAAAGAGGCAGTCCAGTGCGTGCAGGAGCTGGCCTCACCCTCCTTGCTCTTCATCTTTGTACGGCATGGTGTCGAGTCTACGCTGGAGCGCAGTGCCATTGCTCGTGAGCATATGGGGCAGCTGCTGCACCAGCTGCTCTGTGCTGGGCATCTGTCTACTGCTCAGTACTACCAAGGGTTGTATGAAATCTTGGAATTGGCTGAGGACATGGAAATTGACATCCCCCACGTGTGGCTCTACCTAGCGGAACTGGTAACACCCATTCTGCAGGAAGGTGGGGTGCCCATGGGGGAGCTGTTCAGGGAGATTACAAAGCCTCTGAGACCGTTGGGCAAAGCTGCTTCCCTGTTGCTGGAGATCCTGGGCCTCCTGTGCAAAAGCATGGGTCCTAAAAAGGTGGGGACGCTGTGGCGAGAAGCCGGGCTTAGCTGGAAGGAATTTCTACCTGAAGGCCAGGACATTGGTGCATTCGTCGCTGAACAGAAGGTGGAGTATACCCTGGGAGAGGAGTCGGAAGCCCCTGGCCAGAGGGCACTCCCCTCCGAGGAGCTGAACAGGCAGCTGGAGAAGCTGCTGAAGGAGGGCAGCAGTAACCAGCGGGTGTTCGACTGGATAGAGGCCAACCTGAGTGAGCAGCAGATAGTATCCAACACGTTAGTTCGAGCCCTCATGACGGCTGTCTGCTATTCTGCAATTATTTTTGAGACTCCCCTCCGAGTGGACGTTGCAGTGCTGAAAGCGCGAGCGAAGCTGCTGCAGAAATACCTGTGTGACGAGCAGAAGGAGCTACAGGCGCTCTACGCCCTCCAGGCCCTTGTAGTGACCTTAGAACAGCCTCCCAACCTGCTGCGGATGTTCTTTGACGCACTGTATGACGAGGACGTGGTGAAGGAGGATGCCTTCTACAGTTGGGAGAGTAGCAAGGACCCCGCTGAGCAGCAGGGCAAGGGTGTGGCCCTTAAATCTGTCACAGCCTTCTTCAAGTGGCTCCGTGAAGCAGAGGAGGAGTCT GACCACAACGACTACAAGGATGACGATGACAAG<u>T**AA**</u>**GCTT**

### Human eIF4G (84-1600)

<u>ATG</u>CATCATCATCATCATCATGGATCCCAAGTAATGATGATCCCTTCCCAGATCT CCTACCCAGCCTCCCAGGGGGCCTACTACATCCCTGGACAGGGGCGTTCCACATACGTTGTCCCGACACAGCAGTACCCTGTGCAGCCAGGAGCCCCAGGCTTCTATCCAGGTGCAAGCCCTACAGAATTTGGGACCTACGCTGGCGCCTACTATCCAGCCCAAGGGGTGCAGCAGTTTCCCACTGGCGTGGCCCCCGCCCCAGTTTTGATGAACCAGCCACCCCAGATTGCTCCCAAGAGGGAGCGTAAGACGATCCGAATTCGAGATCCAAACCAAGGAGGAAAGGATATCACAGAGGAGATCATGTCTGGGGCCCGCACTGCCTCCACACCCACCCCTCCCCAGACGGGAGGCGGTCTGGAGCCTCAAGCTAATGGGGAGACGCCCCAGGTTGCTGTCATTGTCCGGCCAGATGACCGGTCACAGGGAGCAATCATTGCTGACCGGCCAGGGCTGCCTGGCCCAGAGCATAGCCCTTCAGAATCCCAGCCTTCGTCGCCTTCTCCGACCCCATCACCATCCCCAGTCTTGGAACCGGGGTCTGAGCCTAATCTCGCAGTCCTCTCTATTCCTGGGGA CACTATGACAACTATACAAATGTCTGTAGAAGAATTAACCCCCATCTCCCGTGAAACTGGGGAGCCATATCGCCTCTCTCCAGAACCCACTCCTCTCGCCGAACCCATACTGGAAGTAGAAGTGACACTTAGCAAACCGGTTCCAGAATCTGAGTTTTCTTCCAGTCCTCTCCAGGCTCCCACCCCTTTGGCATCTCACACAGTGGAAATTCATGAGCCTAATGGCATGGTCCCATCTGAAGATCTGGAACCAGAGGTGGAGTCAAGCCCAGAGCTTGCTCCTCCCCCAGCTTGCCCCTCCGAATCCCCTGTGCCCATTGCTCCAACTGCCCAACCTGAGGAACTGCTCAACGGAGCCCCCTCGCCACCAGCTGTGGACTTAAGCCCAGTCAGTGAGCCAGAGGAGCAGGCCAAGGAGGTGACAGCATCAGTGGCGCCCCCCACCATCCCCTCTGCTACTCCAGCTACGGCTCCTTCAGCTACTTCCCCAGCTCAGGAGGAGGAAATGGAAGAAGAAGAAGAAGAGGAAGAAGGAGAAGCAGGAGAAGCAGGAGAAGCTGAGAGTGAGAAAGGAGGAGAGGAACTGCTCCCCCCAGAGAGTACCCCTATTCCAGCCAACTTGTCTCAGAATTTGGAGGCAGCAGCAGCCACTCAAGTGGCAGTATCTGTGCCAAAGAGGAGACGGAAAATTAAGGAGCTAAATAAGAAGGAGGCTGTTGGAGACCTTCTGGATGCCTTCAAGGAGGCGAACCCGGCAGTACCAGAGGTGGAAAATCAGCCTCCTGCAGGCAGCAATCCAGGCCCAGAGTCTGAGGGCAGTGGTGTGCCCCCACGTCCTGAGGAAGCAGATGAGACCTGGGACTCAAAGGAAGACAAAATTCACAATGCTGAGAACATCCAGCCCGGGGAACAGAAGTATGAATATAAGTCAGATCAGTGGAAGCCTCTAAACCTAGAGGAGAAAAAACGTTACGACCGTGAGTTCCTGCTTGGTTTTCAGTTCATCTTTGCCAGTATGCAGAAGCCAGAGGGATTGCCACATATCAGTGACGTGGTGCTGGACAAGGCCAATAAAACACCACTGCGGCCACTGGATCCCACTAGACTACAAGGCATAAATTGTGGCCCAGACTTCACTCCATCCTTTGCCAACCTTGGCCGGACAACCCTTAGCACCCGTGGGCCCCCAAGGGGTGGGCCAGGTGGGGAGCTGCCCCGTGGGCCGGCTGGCCTGGGACCCCGGCGCTCTCAGCAGGGACCCCGAAAAGAACCACGCAAGATCATTGCCACAGTGTTAATGACCGAAGATATAAAACTGAACAAAGCAGAGAAAGCCTGGAAACCCAGCAGCAAGCGGACGGCGGCTGATAAGGATCGAGGGGAAGAAGATGCTGATGGCAGCAAAACCCAGGACCTATTCCGCAGGGTGCGCTCCATCCTGAATAAACTGACACCCCAGATGTTCCAGCAGCTGATGAAGCAAGTGACGCAGCTGGCCATCGACACCGAGGAACGCCTCAAAGGGGTCATTGACCTCATTTTTGAGAAGGCCATTTCAGAGCCCAACTTCTCTGTGGCCTATGCCAACATGTGCCGCTGCCTCATGGCGCTGAAAGTGCCCACTACGGAAAAGCCAACAGTGACTGTGAACTTCCGAAAGCTGTTGTTGAATCGATGTCAGAAGGAGTTTGAGAAAGACAAAGATGATGATGAGGTTTTTGAGAAGAAGCAAAAAGAGATGGATGAAGCTGCTACGGCAGAGGAACGAGGACGCCTGAAGGAAGAGCTGGAAGAGGCTCGGGACATAGCCCGGCGGCGCTCTTTAGGGAATATCAAGTTTATTGGAGAGTTGTTCAAACTGAAGATGTTAACAGAGGCAATAATGCATGACTGTGTGGTCAAACTGCTTAAGAACCATGATGAAGAGTCCCTTGAGTGCCTTTGTCGTCTGCTCACCACCATTGGCAAAGACCTGGACTTTGAAAAAGCCAAGCCCCGAATGGATCAGTATTTCAACCAGATGGAAAAAATCATTAAAGAAAAGAAGACGTCATCCCGCATCCGCTTTATGCTGCAGGACGTGCTGGATCTGCGAGGG AGCAATTGGGTGCCACGCCGAGGGGATCAGGGTCCCAAGACCATTGACCAGATCCATAAGGAGGCTGAGATGGAAGAACATCGAGAGCACATCAAAGTGCAGCAGCTCATGGCCAAGGGCAGTGACAAGCGTCGGGGCGGTCCTCCAGGCCCTCCCATCAGCCGTGGACTTCCCCTTGTGGATGATGGTGGCTGGAACACAGTTCCCATCAGCAAAGGTAGCCGCCCCATTGACACCTCACGACTCACCAAGATCACCAAGCCTGGCTCCATCGATTCTAACAACCAGCTCTTTGCACCTGGAGGGCGACTGAGCTGGGGCAAGGGCAGCAGCGGAGGCTCAGGAGCCAAGCCCTCAGACGCAGCATCAGAAGCTGCTCGCCCAGCTACTAGTACTTTGAATCGCTTCTCAGCCCTTCAACAAGCGGTACCCACAGAAAGCACAGATAATAGACGTGTGGTGCAGAGGAGTAGCTTGAGCCGAGAACGAGGCGAGAAAGCTGGAGACCGAGGAGACCGCCTAGAGCGGAGTGAACGGGGAGGGGACCGTGGGGACCGGCTTGATCGTGCGCGGACACCAGCTACCAAGCGGAGCTTCAGCAAGGAAGTGGAGGAGCGGAGTAGAGAACGGCCCTCCCAGCCTGAGGGGCTGCGCAAGGCAGCTAGCCTCACGGAGGATCGGGACCGTGGGCGGGATGCCGTGAAGCGAGAAGCTGCCCTACCCCCAGTGAGCCCCCTGAAGGCGGCTCTCTCTGAGGAGGAGTTAGAGAAGAAATCCAAGGCTATCATTGAGGAATATCTCCATCTCAATGACATGAAAGAGGCAGTCCAGTGCGTGCAGGAGCTGGCCTCACCCTCCTTGCTCTTCATCTTTGTACGGCATGGTGTCGAGTCTACGCTGGAGCGCAGTGCCATTGCTCGTGAGCATATGGGGCAGCTGCTGCACCAGCTGCTCTGTGCTGGGCATCTGTCTACTGCTCAGTACTACCAAGGGTTGTATGAAATCTTGGAATTGGCTGAGGACATGGAAATTGACATCCCCCACGTGTGGCTCTACCTAGCGGAACTGGTAACACCCATTCTGCAGGAAGGTGGGGTGCCCATGGGGGAGCTGTTCAGGGAGATTACAAAGCCTCTGAGACCGTTGGGCAAAGCTGCTTCCCTGTTGCTGGAGATCCTGGGCCTCCTGTGCAAAAGCATGGGTCCTAAAAAGGTGGGGACGCTGTGGCGAGAAGCCGGGCTTAGCTGGAAGGAATTTCTACCTGAAGGCCAGGACATTGGTGCATTCGTCGCTGAACAGAAGGTGGAGTATACCCTGGGAGAGGAGTCGGAAGCCCCTGGCCAGAGGGCACTCCCCTCCGAGGAGCTGAACAGGCAGCTGGAGAAGCTGCTGAAGGAGGGCAGCAGTAACCAGCGGGTGTTCGACTGGATAGAGGCCAACCTGAGTGAGCAGCAGATAGTATCCAACACGTTAGTTCGAGCCCTCATGACGGCTGTCTGCTATTCTGCAATTATTTTTGAGACTCCCCTCCGAGTGGACGTTGCAGTGCTGAAAGCGCGAGCGAAGCTGCTGCAGAAATACCTGTGTGACGAGCAGAAGGAGCTACAGGCGCTCTACGCCCTCCAGGCCCTTGTAGTGACCTTAGAACAGCCTCCCAACCTGCTGCGGATGTTCTTTGACGCACTGTATGACGAGGACGTGGTGAAGGAGGATGCCTTCTACAGTTGGGAGAGTAGCAAGGACCCCGCTGAGCAGCAGGGCAAGGGTGTGGCCCTTAAATCTGTCACAGCCTTCTTCAAGTGGCTCCGTGAAGCAGAGGAGGAGTCTGACCA CAACGGATCCCGGGCTGACTACAAGGATGACGATGACAAG<u>TAA</u>

